# Temperature-dependent fasciation mutants connect mitochondrial RNA processing to control of lateral root morphogenesis

**DOI:** 10.1101/2020.06.09.141382

**Authors:** Kurataka Otsuka, Akihito Mamiya, Mineko Konishi, Mamoru Nozaki, Atsuko Kinoshita, Hiroaki Tamaki, Masaki Arita, Masato Saito, Kayoko Yamamoto, Takushi Hachiya, Ko Noguchi, Takashi Ueda, Yusuke Yagi, Takehito Kobayashi, Takahiro Nakamura, Yasushi Sato, Takashi Hirayama, Munetaka Sugiyama

## Abstract

Although mechanisms that activate organogenesis in plants are well established, much less is known about the subsequent fine-tuning of cell proliferation, which is crucial for creating properly structured and sized organs. Here we show, through analysis of temperature-dependent fasciation (TDF) mutants of Arabidopsis, *root redifferentiation defective 1* (*rrd1*), *rrd2*, and *root initiation defective 4* (*rid4*), that mitochondrial RNA processing is required for limiting cell division during early lateral root (LR) organogenesis. These mutants formed abnormally broadened (i.e., fasciated) LRs under high-temperature conditions due to excessive cell division. All TDF proteins localized to mitochondria, where they were found to participate in RNA processing: RRD1 in mRNA deadenylation, and RRD2 and RID4 in mRNA editing. Further analysis suggested that LR fasciation in the TDF mutants is triggered by reactive oxygen species generation caused by defective mitochondrial respiration. Our findings provide novel clues for the physiological significance of mitochondrial activities in plant organogenesis.

## Introduction

Plants elaborate their architecture by continuously developing new organs, such as leaves, floral organs, axillary stems, and lateral roots (LRs). Organogenesis begins with the local activation of cell proliferation in the plant body. In the following stages, proliferation is restricted to certain areas, which is essential for the formation of properly sized and structured organs. However, the molecular underpinnings of such regulation remain mostly unknown.

LRs serve as building blocks of the root system architecture, and are crucial for the uptake and transport of water and minerals. The first visible step of LR formation occurs within the parent root, where a few cells start to divide, comprising the LR primordium. The LR primordium grows and eventually emerges out of the parent root to form a new LR [1]. This process has been described in detail in the model plant *Arabidopsis thaliana* (Arabidopsis), rendering it one of the most ideal systems to study the molecular mechanisms of organ development [3,4]. In Arabidopsis, a small number of cells in a few adjacent files of the xylem pole pericycle layer, termed LR founder cells, re-enter the cell cycle and first divide in the anticlinal (perpendicular to the parental root axis) orientation (Fig. 1B) [3,4]. The local accumulation of the phytohormone auxin is critical for LR initiation, driving LR founder cell identity acquisition and division via the degradation of the SOLITARY ROOT (SLR/IAA14) repressor, thus activating the expression of down-stream genes mediated by the AUXIN RESPONSE FACTORS ARF7 and ARF19 [5]. However, much less is understood about the coordinated periclinal (parallel to the surface of the root) and anticlinal divisions that subsequently take place. In particular, the manner in which cell proliferation becomes confined to the central zone of the primordium, giving rise to the dome-shaped structure, largely remains a mystery [1], although the requirement of several factors, such as polar auxin transport [6,7], control of auxin response [8], a few peptide hormones [9,10], transcription factors [11,12], symplastic connectivity [13], epigenetic gene regulation [14], and mechanical interaction with the overlaying tissue [15], has been revealed.

**Fig. 1.**
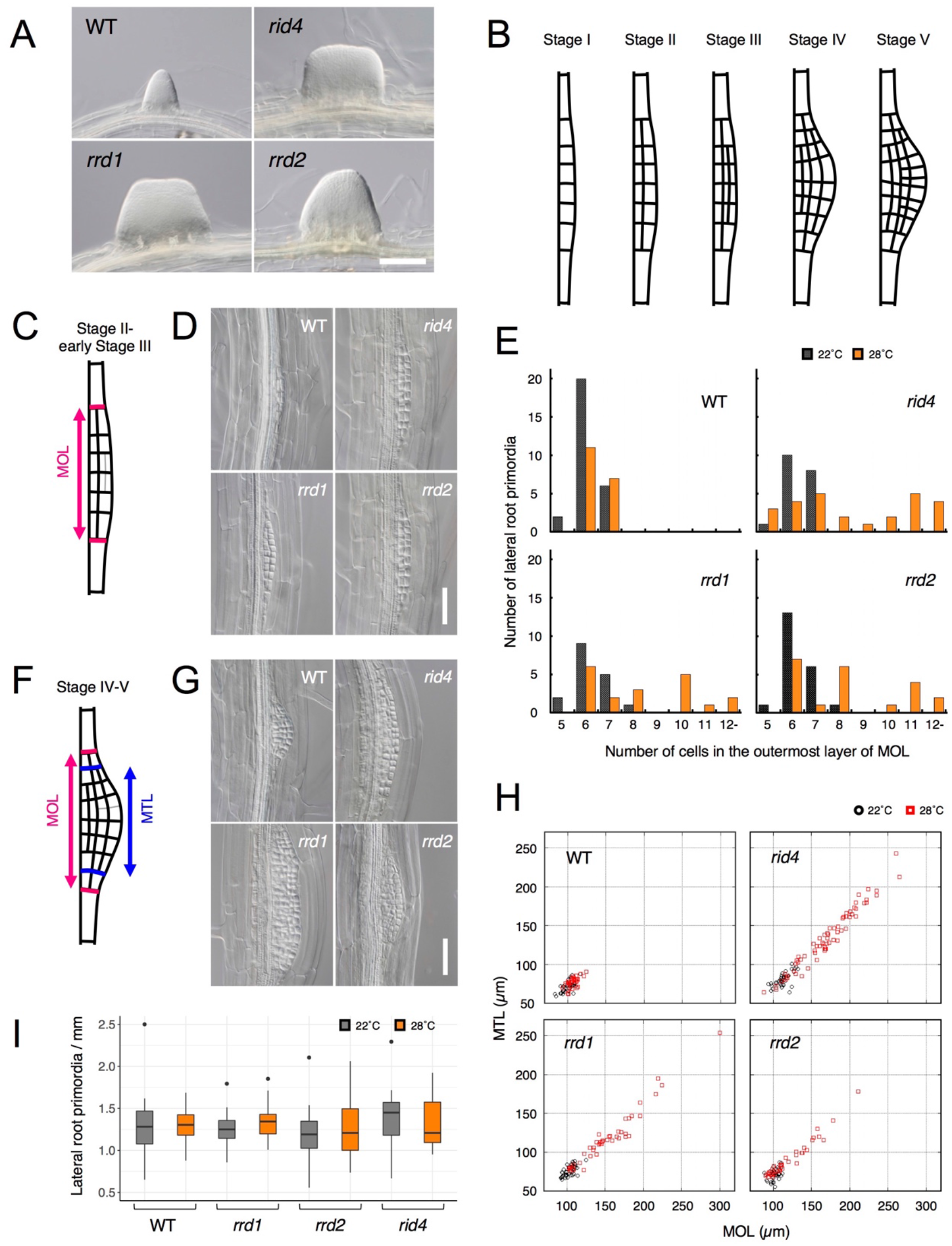
Effects of the TDF mutations on the early stages of LR development. **(A)** Fasciated LRs formed at 28°C in the TDF mutant explants vs. a normal root on the wild-type (WT) explant after 6 days of culture. **(B)** Schematic representation of LR development (stages I–V). **(C)** Schematic image of a primordium at stage II. The area consisting of more than one cell layer (MOL) is delimited by red lines. **(D)** Stage II primordia formed at 28°C in WT and TDF mutant explants. **(E)** Effects of the TDF mutations on the number of cells in the outermost layer of the MOL area of stage II primordia at 22°C (black) and 28°C (orange). N = 17–28. **(F)** Schematic image of a primordium at the transition from stage IV to stage V. The areas consisting of MOL and more than two cell layers (MTL) are delimited by red lines and blue lines, respectively. **(G)** Stage IV–V primordia formed at 28°C in WT explants and TDF mutant explants. **(H)** Scatterplot of the effect of the TDF mutations on the width of the MTL vs. the width of the MOL areas at 22°C (black) and 28°C (red). N = 31–66. **(I)** LR densities in the WT explants and TDF mutant explants cultured at 22°C or 28°C (including all developmental stages; median, 25%–75% quantile, N = 21–29, *P* > 0.3, Kruskal-Wallis test). Scale bars, 100 μm (**A**), 50 μm (**D, G**).

*root redifferentiation defective 1* (*rrd1*), *rrd2*, and *root initiation defective 4* (*rid4*) are temperature-sensitive mutants of Arabidopsis that were originally isolated by us via screening using adventitious root (AR) formation from hypocotyl tissue segments as an index phenotype [16,17]. In addition to AR formation, other aspects of development, such as seedling growth and callus formation, were affected by high-temperature conditions [16,17]. Most notable among these aspects was their LR phenotype, in which abnormally broadened (i.e., fasciated) LRs were formed at 28°C (non-permissive temperature), but not at 22°C (permissive temperature), in a tissue culture setting; thus, we termed the three mutants as temperature-dependent fasciation (TDF) mutants [18]. It was later revealed that the early stages of LR development are likely affected in the TDF mutants, and that the fasciated LRs exhibit exclusive enlargement of inner tissues [18], suggesting that the genes responsible for the TDF mutations (TDF genes) encode negative regulators of proliferation that are important for the size restriction of the central zone during the formation of early stage LR primordia; however, their molecular identity has remained elusive.

Plant cells have gene expression systems in mitochondria and plastids in addition to the nucleus. Although organelle gene expression is typically associated with organelle-specific functions, it might also be involved in higher-order physiological activities including the regulation of organogenesis. Mitochondria are considered the “powerhouses” of the cell, as they supply the energy that is necessary for cellular activities. In comparison with other eukaryotes, RNA metabolism in mitochondria is particularly complex in plants, and entails numerous nuclearly encoded RNA-binding proteins [2]. Given the relaxed nature of transcription, post-transcriptional processing, such as RNA editing, splicing, maturation of transcript ends and RNA degradation, are known to play predominant roles in shaping the plant mitochondrial transcriptome [2]. Many factors that participate in plant mitochondrial RNA processing have been identified; however, the implications of their role in regulating plant organ development remain unclear [2].

Herein, we report a detailed analysis of the TDF mutants. We found that LR fasciation in the TDF mutants was caused by excessive cell division in the early stages of LR formation. Next, we identified all three TDF genes as encoding nuclearly encoded mitochondrial RNA processing factors. Analysis of mitochondrial RNA demonstrated that RRD1 is involved in the removal of poly(A) tails, and that both RRD*2* and RID4 are RNA editing factors. Defective protein composition of the mitochondrial electron transport chain was found in *rrd2* and *rid4*. Phenocopying of the TDF mutants by mitochondrial respiratory inhibition and reactive oxygen species (ROS) induction, together with its reversal by ROS scavenging, suggested that ROS generation resulting from impaired RNA processing is the primary cause of the excessive cell division observed during early LR development in the TDF mutants. Our discovery shed light on a new aspect of mitochondrial RNA processing that is relevant in the control of plant organogenesis.

## Results

### Effects of the TDF mutations on LR formation

To gain insight into fasciated LR formation in the TDF mutants, a detailed investigation was carried out using the semi-synchronous LR induction system [19], in which nearly de novo LR formation is induced from root explants of young seedlings upon culture in auxin-containing root inducing medium (RIM). In this system, a 6-day culture of TDF mutant explants results in high rates of LR fasciation at 28°C (non-permissive temperature) (Fig. 1A), but not at 22°C (permissive temperature) [18]. To determine the stage of LR formation at which developmental abnormalities occur in the TDF mutants, LR primordia from earlier time points were examined. In Arabidopsis, LR formation begins with anticlinal cell divisions in the xylem pole pericycle cell file, producing an array of short cells flanked by longer cells, which serve as the origin of the LR primordium (stage I; Fig. 1B) [3,4]. This is followed by periclinal divisions throughout the primordium, with the exception of the flanking cells in some occasions, creating two cell layers (stage II; Fig. 1B). Subsequent periclinal cell divisions take place in the central zone of the primordium, producing the third cell layer (stage III), followed by the fourth cell layer (stage IV; Fig.1B). Additional anticlinal cell division, together with cell expansion at the innermost cell layer, gives rise to a dome-shaped primordium (stage V; Fig. 1B). The comparison of the number of cells within the area consisting of more than one layer (MOL) [11] between stages II and III, revealed that all TDF mutants showed an increase in this parameter in a temperature-dependent manner (Fig. 1, C to E). The same trend was observed in primordia at stage IV and V, for which the widths of the MOL and more than three layer (MTL) [11] areas were quantified (Fig. 1, F to H). These results showed that TDF mutants exhibit excessive cell division in the initial steps of LR development, namely as early as stage II, and indicate that the increase in the number of cells along the lateral axis of the primordium induces the expansion of its central zone, giving rise to an abnormally broadened and flat-shaped LR. As there was no significant increase in LR density (Fig. 1I; Kruskal-Wallis test, P > 0.3), LR fasciation in the TDF mutants seems to be the result of the expansion of individual primordia, as opposed to the fusion of multiple primordia because of overcrowding that is observed in some other mutants [9,13].

### Positional cloning and expression analysis of the TDF genes

To clone the TDF genes, we mapped the mutated loci in the TDF mutants based on the temperature-sensitive AR formation phenotype, which originally led to the isolation of the mutants (fig. S1) [16,17]. The candidate genes identified by sequencing the mapped regions were confirmed either by a complementation test (*RRD1* and *RID4*; fig. S2, A and E) or an allelism test (*RRD2*; fig. S2, B to D). This resulted in the identification of *RRD1* as At3g25430, which encodes a poly(A)-specific ribonuclease (PARN)-like protein, and *RRD2* and *RID4* as At1g32415 and At2g33680, respectively, both of which encode a pentatricopeptide repeat (PPR) protein belonging to the PLS subfamily (Fig. 2A). At1g32415 has previously been reported as the gene responsible for the *cell wall maintainer 2* (*cwm2*) mutation [20]; thus, we will refer to it as *RRD2/CWM2* henceforth. *rrd1, rrd2*, and *rid4-1* are all nonsense mutations (Fig. 2A). The *rrd1* mutation results in an 89-amino-acid C-terminal truncation of the 618-amino-acid RRD1 protein; the mutant protein may be partially or conditionally functional. As the *rrd2* and *rid4* mutations create a stop codon close to the start codon (Fig. 2A), they are likely to eliminate gene function. Later in our study, another mutant harboring a mutation in the *RID4* gene was isolated and designated *rid4-2* (Fig. 2A and fig. S3). *rid4-2* exhibited LR fasciation as well as retarded seedling growth at high-temperature conditions, similar to *rid4-1* (fig. S3, A and B). The *rid4-2* mutation is a missense mutation that gives rise to a single amino acid substitution (G137R) (Fig. 2A and fig. S3D), presumably causing a partial reduction of gene function.

**Fig. 2.**
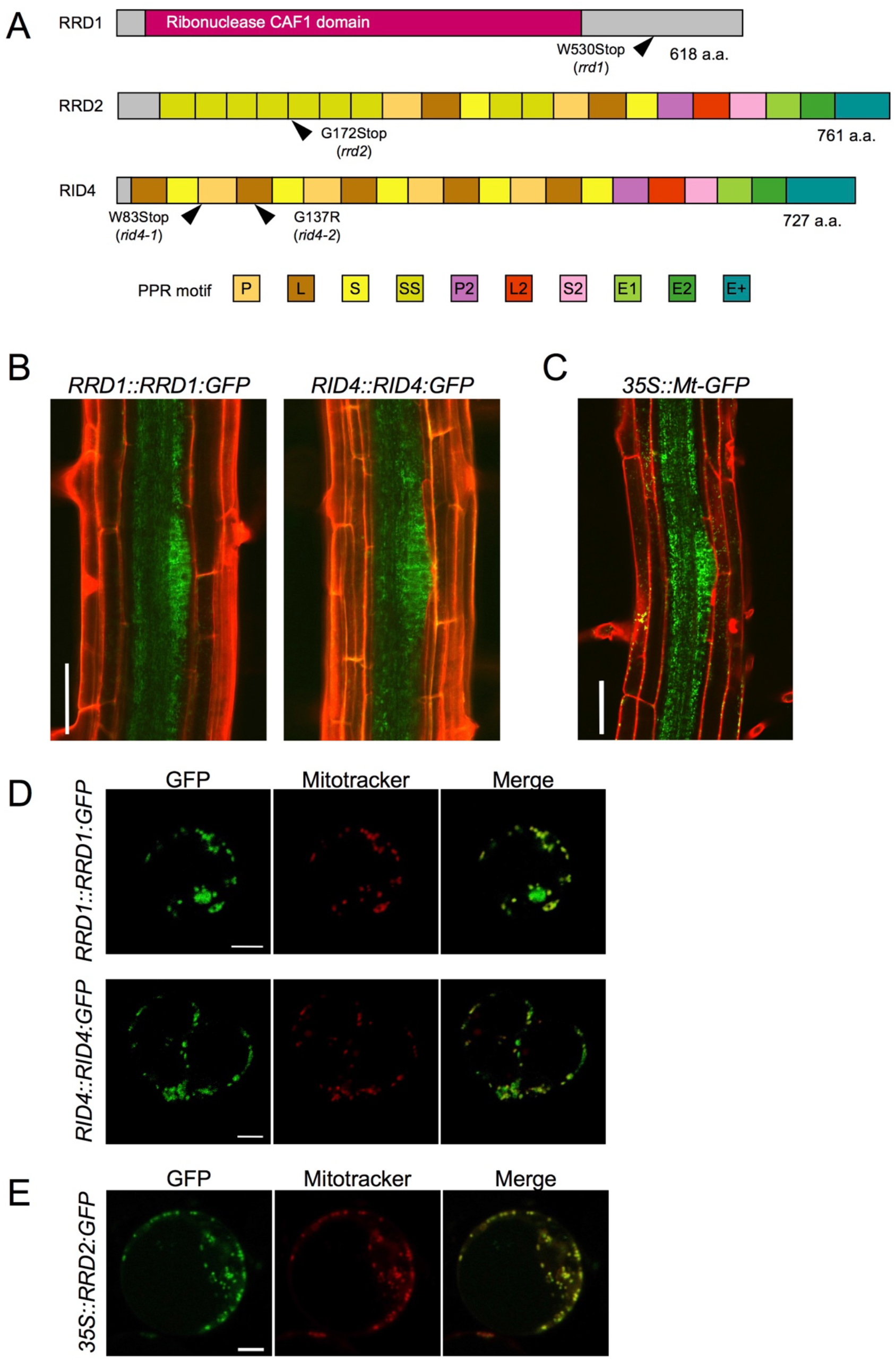
Tissue-specific expression and subcellular localization of the TDF proteins. **(A)** Structures of the RRD1, RRD2, and RID4 proteins. **(B and C)** Expression of *RRD1::RRD1:GFP* (**B**, left), *RID4::RID4:GFP* (**B**, right), and *35S::Mt-GFP* (**C**) at stage II of LR primordium development. Propidium iodide was used as a red counterstain. **(D and E)** Expression of *RRD1::RRD1:GFP* (**D**, upper panels), *RID4::RID4:GFP* (**D**, lower panels), and *35S::RRD2:GFP* (**E**) in callus-derived protoplasts. Mitochondria were labeled with MitoTracker Orange. Scale bars, 50 μm (**B and C**) and 5 μm (**D and E**).

GFP reporter studies were carried out to elucidate the expression patterns of the TDF genes. For *RRD1* and *RID4*, genomic constructs encompassing the promoter region to the end of the protein-coding sequence (*RRD1::RRD1:GFP* and *RID4::RID4:GFP*) were generated and introduced into *rrd1* and *rid4-1*, respectively. The suppression of the mutant AR phenotype demonstrated the functionality of the reporter genes (fig. S4, A and B). For both *RRD1* and *RID4*, strong GFP expression was mostly confined to apical meristems and LR primordia in the root system and slightly and much weaker expressions were detected in the stele and cortex/epidermis tissues, respectively (Fig. 2B, and fig. S4C). This resembled the *35S::Mt-GFP* line, which expresses mitochondria-targeted GFP under the constitutive active cauliflower mosaic virus (CaMV) 35S promoter (Fig. 2C). At the subcellular level, fluorescence from the GFP-fusion proteins appeared punctate or granulated and was largely overlapped with signals from the mitochondrion-specific dye MitoTracker Orange, demonstrating that the majority of RRD1 and RID4 proteins are localized to mitochondria (Fig. 2D). Although the tissue-level investigation of *RRD2/CWM2* expression was unsuccessful because of the undetectable levels of the signals of *RRD2::RRD2:GFP*, mitochondrial localization was also confirmed for RRD2 by studying transient expression under the 35S promoter (Fig. 2E). Together, these data showed that the TDF genes *RRD1, RRD2/CWM2*, and *RID4* encode putative RNA processing factors that localize to mitochondria.

### Analysis of the role of RRD1 in poly(A) degradation of mitochondrial mRNAs

PARN belongs to the DEDD superfamily of deadenylases [21]. Recent human and animal studies have led to an increased appreciation of its participation in the maturation process of a wide variety of noncoding RNAs [22]. In plants, however, PARN plays a distinct role in the removal of the poly(A) tails of mitochondrial mRNA [23–25]. Given the sequence similarity to PARN and its mitochondrial localization, we hypothesized that RRD1 is also involved in regulating the poly(A) status of mitochondrial mRNA. To test this possibility, we first performed a microarray analysis of poly(A)^+^ RNAs prepared from wild-type and *rrd1* explants that had been induced to form LRs at 28°C, and found a substantial increase in mitochondria-encoded poly(A)^+^ transcripts in *rrd1* explants (Fig. 3A, and fig. S5, A to C). As the majority of plant mitochondrial transcripts normally lack poly(A) tails, presumably because of swift removal after its addition [26], we suspected that the apparent sharp increase in mitochondrial transcript level might be ascribed to defective poly(A) tail removal, rather than increased transcription. In fact, a comparative analysis of polyadenylated and total RNA levels via quantitative reverse transcription polymerase chain reaction (qRT-PCR) revealed a selective increase in polyadenylated transcripts (Fig. 3B). Furthermore, a circularized RNA (CR)-RT PCR analysis [27] of the *cytochrome oxidase subunit 1* (*cox1*) mRNA was performed to study its 3’ extremity, and revealed a marked increase in the polyadenylated to non-polyadenylated ratio in *rrd1* compared with the wild-type plant (Fig. 3C). In addition, a poly(A) test assay by rapid amplification of cDNA ends (RACE-PAT) [28] showed that polyadenylated transcript levels were increased at higher temperature in *rrd1* (Fig. 3D). Taken together, these results demonstrated that RRD1 is involved in poly(A) tail removal in mitochondrial mRNAs, and that, in *rrd1*, polyadenylated mitochondrial transcripts accumulate in a temperature-dependent manner.

**Fig. 3.**
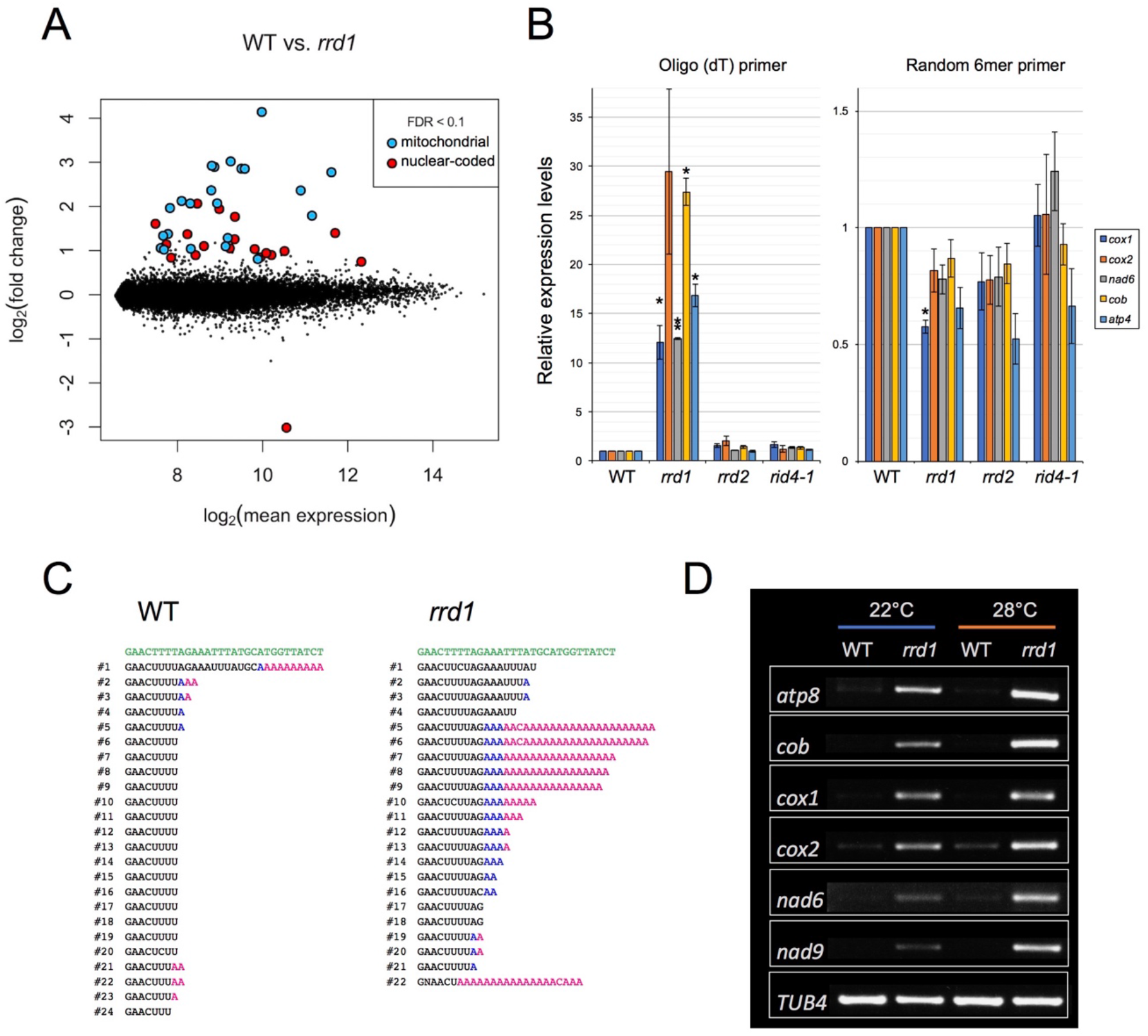
Accumulation of polyadenylated mitochondrial transcripts in *rrd1*. **(A)** MA plot for the microarray analysis of poly(A)^+^ transcripts of *rrd1* vs. wild-type (WT) explants in which LRs were induced at 28°C for 12 hours. **(B)** qRT–PCR analysis of explants in which LRs were induced at 28°C for 12 hours. The total and polyadenylated transcript levels are shown for *cytochrome oxidase subunit 1* (*cox1*), *cox2, NADH dehydrogenase subunit 6* (*nad6*), *apocytochrome B* (*cob*), and *ATP synthase subunit 4* (*atp4*) (mean ± s.d., N = 3, \**P* < 0.05, \*\**P* < 0.01, 1 sample *t* test with Benjamini-Hochberg correction). **(C)** Analysis of the 3’ end of the *cox1* mRNA by CR–RT PCR. mRNAs were prepared from WT and *rrd1* seedlings that were first grown at 22°C for 7 days, and then at 28°C for 2 days. The genomic sequence of *cox1* is shown in green. **(D)** RACE-PAT assay showing the accumulation of polyadenylated transcripts of *atp8, cob, cox1, cox2, nad6, nad9*, and *TUB4*. mRNAs were prepared from explants in which LRs were induced at 22°C or 28°C for 12 hours.

Next, we investigated whether the RRD1 protein itself has deadenylation activity. In previous studies, this possibility was excluded because, in contrast to canonical PARNs (including AtPARN/AHG2), RRD1 lacks three out of the four amino acids that are essential for its function as a deadenylase [29]. In our assay, as expected, the recombinant RRD1 protein did not show any activity in the conditions effective for human PARN (fig. S5, D and E). We concluded that the RRD1 protein alone does not have deadenylase activity.

To assess the effects of the observed accumulation of poly(A)^+^ mitochondrial transcripts in *rrd1*, we introduced the *ahg2-1* suppressor *1* (*ags1*) mutation into *rrd1. ags1* is a mutation of a mitochondrion-localized poly(A) polymerase (PAP), AGS1, which was originally identified based on its ability to counteract AtPARN/AHG2 loss of function [23]. A substantial decrease in mitochondrial poly(A)^+^ transcript levels was observed in the *rrd1 ags1* double mutant compared with the *rrd1 AGS1* control (Fig. 4A). Moreover, *rrd1* phenotypes, such as temperature-dependent LR fasciation and seedling growth retardation, were significantly alleviated (Fig. 4, B and C). These results indicate that the accumulation of poly(A)^+^ mitochondrial transcripts is the primary cause of the *rrd1* phenotype.

**Fig. 4.**
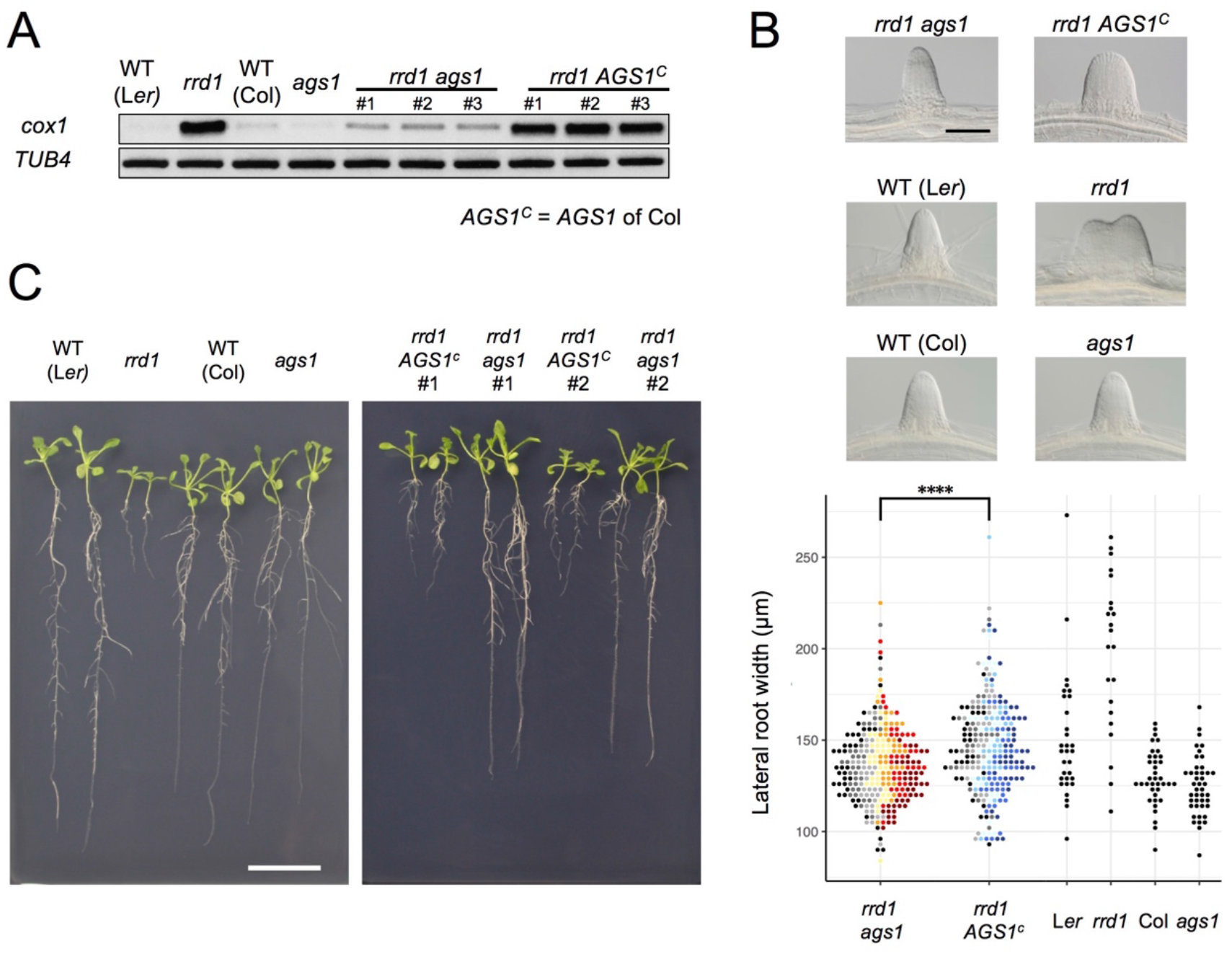
Effects of *ags1* on the phenotypes of *rrd1*. **(A)** RACE-PAT assay showing the accumulation of polyadenylated transcripts of *cox1* and *TUB4. rrd1* mutant strains harboring either *ags1* or *AGS1*^*c*^ (*AGS1* of Col background) were obtained by *rrd1* (L*er* background) *× ags1* (Col background) and *rrd1 ×* Col crosses, respectively. mRNAs were prepared from seedlings that were first grown at 22°C for 5 days, and then at 28°C for 3 days. **(B)** Representative images of LRs formed at 28°C after 6 days of culture (upper panels). The basal width of the LRs that were formed in this way was scored (lower panel, N = 115–116 for *rrd1 ags1*, and *rrd1 AGS1*^*c*^, N = 22–43 for others, \*\*\*\**P* < 10^−4^, Mann– Whitney–Wilcoxon test with Bonferroni correction). For *rrd1 ags1* and *rrd1 AGS1*^*c*^, data were gathered from 7 strains, which are shown by different colors. **(C)** Seedlings grown at 28°C for 13 days on gellan gum plates. Scale bars, 100 μm (**B**) and 2 cm (**C**).

### Analysis of the roles of RRD2 and RID4 in mitochondrial mRNA editing

PPR proteins are known for their role in regulating various aspects of organellar post-transcriptional gene expression, such as RNA stabilization, RNA cleavage, RNA splicing, RNA editing, and translation [2,30]. They are characterized by the tandem assembly of degenerate protein motifs of about 35 amino acids, termed PPR motifs [30]. The PPR motifs allow PPR proteins to recognize specific sites of single-stranded RNAs through a one-motif to one-base interaction [30]. The PPR protein family has undergone a remarkable expansion in land plants, representing one of the largest protein families thereof [30]. RRD2 and RID4 belong to the PLS-class of PPR proteins, most of which have been reported as being C-to-U RNA editing factors [31]. The PLS class PPR proteins contain three types of PPR motifs, the P motif (normally 35 a. a. in length), the L motif (35–36 a. a. (long)) and the S motif (31 a. a. (short)), in contrast to the P-class PPR proteins, which only contain P motifs [30,32]. Considering their localization to mitochondria (Fig. 2, D and E), we speculated on the involvement of RRD2 and RID4 in the editing of mitochondrial RNA. A comprehensive sequence analysis of previously reported RNA editing sites using cDNA prepared from explants induced to form LRs at 28°C revealed an almost complete abolishment of C-to-U editing at two sites (*cytochrome c biogenesis protein 2* (*ccb2*)-71C and *ccb3*-575C) in *rrd2* and at six sites (*ATP synthase subunit 4* (*atp4*)-395C, *ribosomal protein l5* (*rpl5*)-58C, *rpl5*-59C, *rps3*-1344C, *rps4*-77C, and *rps4*-332C) in *rid4* (Fig. 5A, fig. S7). The identification of *ccb3*-575C as an RRD2/CWM2 editing site was in agreement with a previous study of *cwm2* [20]. Editing was also completely abolished in these sites at 22°C (fig. S8A). RID4 editing sites showed incomplete editing in *rid4-2*, implying a partial loss of function in this mutant (fig. S7). Significant identity was found among the 5*’* upstream sequences of the editing sites that were affected in each mutant (fig. S8B), further suggesting that RRD2 and RID4 participate in the editing of these sites via direct contact.

**Fig. 5.**
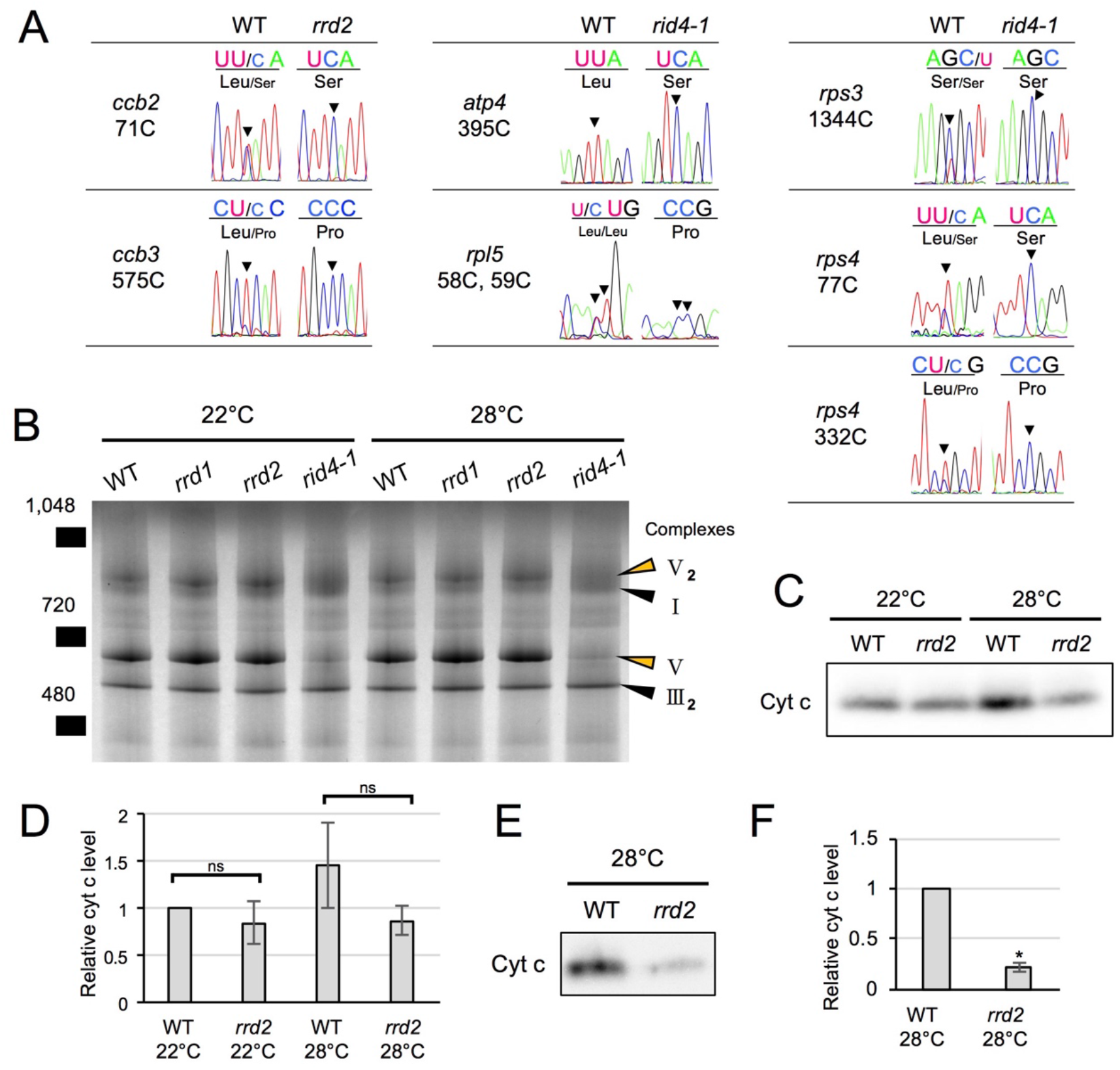
Effects of *rrd2* and *rid4* on mitochondrial mRNA editing and protein synthesis. **(A)** Sequencing analysis of mitochondrial mRNA editing in explants in which LRs were induced at 28°C for 12 hours. **(B)** BN-PAGE analysis of mitochondrial protein complexes. Mitochondria were extracted from seed-derived liquid-cultured callus that were first incubated at 22°C for 20 days, and then at 22°C or 28°C for an additional 3 days. **(C and D)** Immunoblot analysis of cyt c. Mitochondria were extracted in the same conditions as in (B). The results of the densitometry analysis are shown in (**D**) (N = 3, mean ± s.d.). **(E and F)** Immunoblot analysis of cyt c using mitochondria extracted from callus that were cultured first at 22°C for 14 days, and then at 28°C for 7 days. The results of the densitometry analysis are shown in (**F**) (N = 2, mean ± s.d., \*\**P* < 0.01, Welch’s *t* test)

In addition, all editing sites of *ccb3*, with the exception of those that were unedited in the wild type, showed declining levels of RNA editing in both *rrd2* and *rid4* (fig. S7). However, these sites were not considered as targets of RRD2 and RID4 for the following reasons. These sites were incompletely edited, even in the wild type, as opposed to most other sites (fig. S7), suggesting that their editing is relatively slow and highly susceptible to fluctuations in the kinetic balance between editing and transcription. Moreover, editing at these sites was almost unaffected at 22°C (fig. S8C) and was only partially inhibited at 28°C in *rrd2* and *rid4* (fig. S7), even though these mutants are assumed to have lost the function of the corresponding genes completely. *ccb3*-624C was also not regarded as a target site, despite the complete absence of editing in both *rrd2* and *rid4*, as it was more likely due to originally low levels of editing compared with other sites in *ccb3* (fig. S7). This view was reinforced by the lack of similarity in the upstream sequence between *ccb3*-624C and the other editing sites that were strongly affected by the *rrd2* and *rid4* mutations (fig. S8B).

Next, to investigate the effects of losses of function of RRD2/CWM2 and RID4 on mitochondrial protein composition, we performed a blue-native (BN)-PAGE analysis of mitochondrial extracts prepared from seed-derived callus cultured for 3 days at 22°C or 28°C after a 20-day 22°C incubation period. This revealed a substantial loss of complex V (ATP synthase complex) in *rid4* at both 22°C and 28°C culture conditions (Fig. 5B), likely caused by defective mRNA editing of *atp4* (Fig. 5A), which is a component of this protein complex. No noticeable differences were found in *rrd1* and *rrd2*. Because *ccb2* and *ccb3*, the two mitochondrial genes that are targeted by RRD2/CWM2, are related to cytochrome c (cyt c) maturation [33], we quantified cyt c levels in *rrd2*. Cyt c levels on a per mitochondrial protein basis were decreased in *rrd2* callus cultured at 28°C for 3 days (Fig. 5, C and D) in two out of three cultures, although the difference was not significant when all three results were included. This decrease in cyt c levels in *rrd2* was in accordance with a previous analysis of *cwm2* [20]. At 22°C, however, no significant difference was observed between *rrd2* and the wild type. Furthermore, we found that the difference in cyt c levels was more pronounced after longer periods of culture at 28°C (Fig. 5, E and F). These results indicate that, in *rrd2*, cyt c maturation activity was affected to a greater extent at higher temperatures, at least in callus, which possesses root-tissue-like properties, possibly explaining the temperature-dependent nature of its phenotype. The data reported above demonstrated that, in both *rrd2* and *rid4*, the production of certain components of the mitochondrial electron transport chain is hampered by defective mRNA editing.

### Effects of defective mitochondrial respiration on LR formation

Based on the results obtained for *rrd1, rrd2*, and *rid4*, we speculated that there might be a relationship between mitochondrial electron transport and cell division control during LR morphogenesis. In fact, the induction of LRs from wild-type explants in the presence of rotenone (complex I inhibitor), antimycin A (complex III inhibitor), or oligomycin (complex V inhibitor) led to LR fasciation, providing evidence that electron transport chain defects are the cause of the TDF LR phenotype (Fig. 6, A to D). To further investigate the underlying molecular pathway, we next asked whether either reduced ATP synthesis, or ROS generation, phenomena that are commonly associated with defective mitochondrial respiration might be involved. We found that the respiratory uncoupler carbonylcyanide m-chlorophenyl-hydrazone (CCCP) did not increase LR width (Fig. 6E), although LR growth inhibition was observed in a dose-dependent manner (Fig. 6F), whereas the ROS inducer paraquat (PQ) triggered a significant fasciation of LRs (Fig. 6, G and H). Furthermore, the application of the ROS scavenger ascorbate resulted in a reversal of the LR broadening induced by PQ treatment (Fig. 6, G and H). The same effect was observed against the *rid4-2* mutation. These data suggest that the increase in the levels of ROS, but not the decrease in the levels of ATP, acts downstream of defective mitochondrial respiration to promote excessive cell division during LR development in the TDF mutants.

**Fig. 6.**
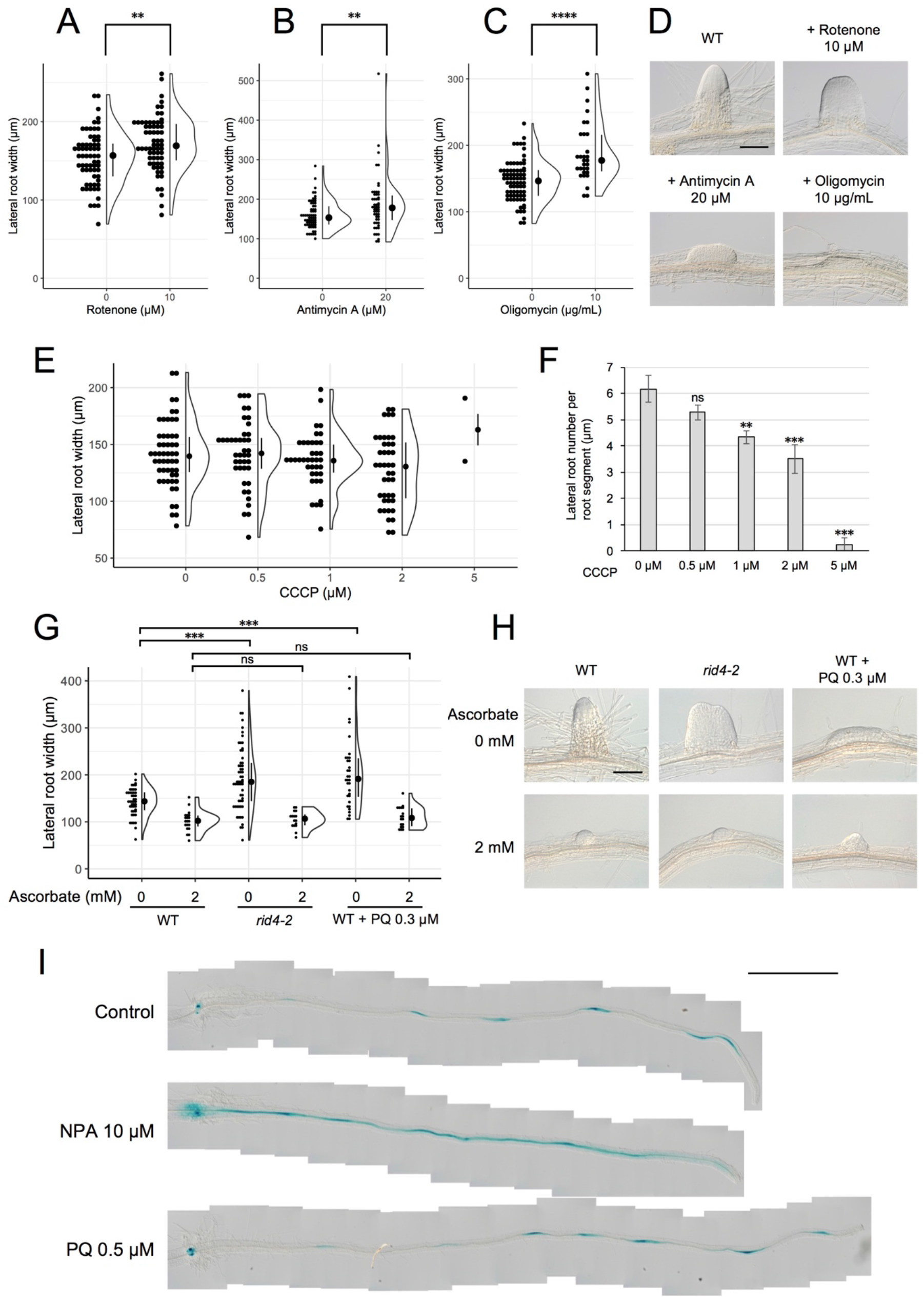
Formation of fasciated LRs after treatment with chemicals that inhibit mitochondrial respiration or induce ROS. **(A–D)** LRs were induced at 28°C from the wild-type (WT) plant in the presence of rotenone (**A**), antimycin A (**B**), or oligomycin (**C**), and the basal width of the LRs that were formed was scored after 6 days in culture (median, 25%–75% quantile, N = 30–76, \*\**P* < 0.01, \*\*\*\**P* < 0.0001, Mann–Whitney–Wilcoxon test). Typical LRs that were formed in each treatment are shown in (**D**). **(E and F)** LRs were induced from the WT plant in the presence of CCCP. The basal width of LRs (**E**, median, 25%–75% quantile, N = 2–53, *P* > 0.1, Kruskal-Wallis test) and the number of LRs per segment (**F**, Number of segments = 12, \*\**P* < 0.01, \*\*\**P* < 0.001, Dunnett’s test) were scored on the 6^th^ day. **(G and H)** The effects of the application of ascorbate on WT, PQ-treated, or *rid4-2* segments during LR formation. The basal width of the LRs formed was measured on the 6^th^ day of LR induction (**G**, median, 25%–75% quantile, N = 16–58, \*\*\**P* < 0.001, Mann–Whitney–Wilcoxon test with Bonferroni correction). Representative images of LRs in each condition are shown in (**H**). **(I)** *DR5::GUS* expression at 12 hours after LR induction under treatment with NPA or PQ. Scale bars, 100 μm (**D** and **H**) and 1 mm (**I**).

Local gradient formation of auxin is important for LR initiation and the subsequent organization of the LR primordium [5–7]. Strong genetic perturbations of polar auxin transport result in homogeneous proliferation of the pericycle cell layer in large regions of the root upon exogenous auxin treatment. In addition, chemical inhibition of auxin polar transport by naphthylphthalamic acid (NPA) gave rise to broadened LR primordia reminiscent of fasciated LRs of the TDF mutants (fig. S9). These data indicate a role for local auxin gradient formation in restricting proliferative capacity during LR formation. Therefore, we tested whether ROS-induced LR fasciation is mediated by altered auxin patterning in early LR primordia. The examination of the expression pattern of the auxin-responsive β-glucuronidase marker *DR5::GUS* [6–8] at early stages of LR induction, however, did not reveal differences between the control and PQ-treated root segments, whereas treatment with NPA resulted in enhanced expression along the entire root segment (Fig. 6I). This result indicates that ROS-induced LR fasciation is not caused by an impairment in auxin gradient formation.

## Discussion

In the present study, we investigated three TDF mutants of Arabidopsis, *rrd1, rrd2*, and *rid4*, which form fasciated LRs at high temperatures, and identified mutations in previously poorly characterized genes encoding mitochondria-localized proteins as being responsible for the phenotype of these mutants. Our results elucidated the roles of these genes in mitochondrial RNA processing, the construction of the respiratory chain, and in the restrictive control of cell proliferation during LR primordium development.

### Excessive cell division during early primordium development leads to LR fasciation

In the present study, we investigated the formation of fasciated LRs observed at high-temperature conditions in the TDF mutants using the semi-synchronous LR induction system [19]. By measuring the cell number and primordium width, we found that fasciation of LRs is caused by excessive anticlinal cell division, which takes place as early as stage II of LR development (Fig. 1). The lack of increase in LR density (Fig. 1I) suggested that LR fasciation is caused by the expansion of individual primordia, rather than the fusion of multiple primordia, which is the case in some other mutants that form abnormally broadened LRs [9,13]. The data are in agreement with the previous result of the temperature-shift experiment, which demonstrated that the first 48 h following LR induction are critical for LR fasciation in the TDF mutants [18], as stage II to early stage III primordia are formed within this time frame (Fig. 2D) [19]. The previous characterization of the TDF mutants also showed that fasciated LR primordia exhibit specific enlargement of inner root tissues marked by the expression of *SHORT ROOT* (*SHR*), while the number of cell layers outside the SHR-expressing layer is normal [18]. A recent study revealed that the area of SHR expression is first established during stage II, where it is confined to the inner layer of the two-cell layered primordium [4]. In subsequent stages, SHR is expressed in cell files derived from the inner layer, which develop into the stele of the LR [4]. Taken together, these results suggest that differentiation into two cell layers at stage II occurs normally in the TDF mutants, and that the increase in the number of cells observed at stage II consequently leads to the expansion of the area of SHR expression in the inner cell layer during LR fasciation.

### RRD1 functions in poly(A) tail removal in mitochondrial mRNA

PARN is a 3’ exoribonuclease of the DEDD superfamily [21], which shows a strong preference for adenine [21,22]. In plants, PARN is involved in the removal of poly(A) tails from mitochondrial transcripts [23–25]. Here, we identified *RRD1* as a gene encoding a PARN-like protein (Fig. 2A) that resides in mitochondria (Fig. 2, B and C). Further analysis of *rrd1* demonstrated the participation of RRD1 in poly(A) tail degradation of mitochondrial mRNA (Fig. 3). In plant mitochondria, immature 3’ extremities of mRNA, together with irregular RNAs, such as 3’ misprocessed mRNAs, rRNA maturation by-products, and cryptic transcripts, are known to be polyadenylated before they are degraded by mitochondrial polynucleotide phosphorylase (mtPNPase) [26]. In fact, down-regulation of mtPNPase in Arabidopsis results in the accumulation of long preprocessed mRNAs, as well as irregular RNAs, the majority of which are polyadenylated [26]. In *rrd1*, unusually long preprocessed mRNAs do not seem to accumulate, as the size of RACE-PAT assay products (Fig. 3D) corresponded to that of previously reported mature transcript 3’ ends. Total mitochondrial mRNA levels were unelevated in *rrd1* (Fig. 3B), suggesting that RRD1 is not involved in controlling mRNA abundance by promoting their degradation. Whether by-product accumulation takes place in *rrd1* is not clear. However, given its absence in *ahg2* [23], this is unlikely. Based on these considerations, we concluded that RRD1 plays a distinct role from mtPNPase and seems to be specifically involved in 3’ processing of near-matured mRNA.

The mode of action of the RRD1 protein remains to be solved. The absence of three out of the four catalytic amino acids (DEDD) that are essential for ribonuclease activity (fig. S6) [29], together with the apparent lack of deadenylase activity of the recombinant RRD1 protein (fig. S5, D and E), indicated that RRD1 requires additional factors for its participation in poly(A) tail removal.

Failure in the removal of poly(A) tails from mitochondrial transcripts seems to be the primary cause of the *rrd1* phenotype. This is evidenced by the alleviation of the *rrd1* phenotype by the introduction of a mutation of the mitochondria-localized poly(A) polymerase gene *AGS1* (Fig. 4). As most protein-coding genes in the Arabidopsis mitochondrial genome are involved in the biogenesis of the electron transport chain [2], it is likely that mitochondria of *rrd1* carry defects in respiratory activity. However, the exact impact of the altered poly(A) status of mRNAs in mitochondria on electron transport in *rrd1* remains unclear. Unlike the AtPARN/AHG2 loss-of-function mutant *ahg2*, which shows a reduction in complex III levels [23], no significant difference in respiratory chain composition has been detected in *rrd1* to date (Fig. 5B).

### RRD2 and RID4 function in mitochondrial mRNA editing

Our study identified *RRD2* and *RID4* as At1g32415 and At2g33680, respectively, both of which encode a mitochondria-localized PLS-class PPR protein (Fig. 2). At1g32415 had previously been reported as the gene responsible for the *cwm2* mutant [20]. A predominant role for PLS-class PPR proteins in RNA editing has been demonstrated with more than 50 out of a total of approximately 200 of these proteins in Arabidopsis having been identified as C-to-U editing factors of mitochondria or plastid RNA [31]. A comprehensive analysis of mitochondrial RNA editing revealed the abolishment of editing at specific sites in *rrd2* and *rid4* (Fig. 5A and fig. S7). We concluded that both RRD2/CWM2 and RID4 are PLS-class PPR proteins that are involved in mitochondrial mRNA editing.

In *rrd2*, editing at 71C of *ccb2* and 575C of *ccb3* was absent (Fig. 5A). Both *ccb2* (also known as *ccb206, ccmB, ABCI2*, and *AtMg00110*) and *ccb3* (also known as *ccb256, ccmC, ABCI3*, and *AtMg00900*) encode a multisubunit ATP-binding cassette (ABC) protein, which are involved in the maturation of mono hemic c-type cytochromes, the soluble cyt c, and the membrane-bound cyt c_1_ of complex III [33]. Of the two editing sites, *ccb3*-575C was previously reported as a target of *RRD2/CWM2* [20], whereas *ccb2*-71C is a newly discovered target. A decrease in the level of cyt c was detected in *rrd2*, which is consistent with that reported previously for *cwm2* [20]. The data demonstrated the role of *RRD2/CWM2* in cyt c maturation via the RNA editing of cyt c biogenesis factors.

In *rid4*, we observed striking reductions in RNA editing at *atp4*-395C, *rpl5*-58, *rpl5*-59C, *rps3*-1344C, *rps4*-77C, and *rps4*-332C. *atp4* (also known as *orf25, AtMg00640*) encodes the peripheral stalk protein (subunit b) of the mitochondrial ATP synthase complex (complex V) [34]. *rpl5, rps3*, and *rps4* encode mitochondrial ribosome proteins. Analysis of mitochondrial protein complexes showed a dramatic decrease in the level of complex V in *rid4*, probably because of impaired editing of *atp4*-395C. This is similar to the *organelle transcript processing 87* (*otp87*) mutant of Arabidopsis, in which editing of *atp1*-1178C is deficient [35]. These data showed that the formation of complex V could be disrupted by defective RNA editing at a single site of a subunit gene. Considering that the C-to-U editing of the *rps4* transcript at a different site (*rps4*-377) has been shown to affect mitochondrial ribosome assembly in the *growing slowly 1* (*grs1*) mutant [35], it is possible that the *rid4* mutation also has an impact on the mitochondrial ribosome.

Recent advances in the mechanistic understanding of RNA binding by PLS-class PPR proteins have led to the identification of residues at certain positions within the PPR motifs that are important for ribonucleotide recognition [30,31]. By mapping these residues of previously reported RNA-editing PPR proteins to their binding sites, which are located 5’ upstream of the editing sites, the so-called ‘PPR code’ has been elucidated, thus enabling the matching of PPR proteins to their candidate editing targets, and vice versa [31]. According to the recently refined PPR code prediction [31], RID4 was highly ranked as a potential binding protein of *atp4*-395C (18^th^, P = 4.35 *×* 10^−2^), *rpl5*-58C (5^th^, P = 3.04 *×* 10^−2^) and *rps4*-332C (2^nd^, P = 4.06 *×* 10^−3^). Conversely, these sites were among the predicted editing sites of RID4 (P < 0.05) [31]. With regard to RRD2, however, the newly identified binding site (*ccb2*-71C) ranked very low, despite the incorporation of RRD2/CWM2 binding to *ccb3*-575C as learning data for the PPR code prediction [31]. This discrepancy may be related to the unusual arrangement of PPR motifs in RRD2, in which repeats of SS motifs are prevalent, in contrast to canonical PLS-class PPRs, which follow the (P1-L1-S1)_n_-P2-L2-S2 pattern, such as RID4 (Fig. 2A) [32]. Nevertheless, given the similarity between the upstream sequences of editing sites which are severely affected by *rrd2* and *rid4* (fig. S8B), they are likely edited by RRD2 and RID4 via direct interaction. The presented data will contribute to the improvement of PPR protein target estimation.

### The origins of the temperature sensitivity may differ among the TDF mutants

A distinct feature of the TDF phenotype is its exclusive observation at high-temperature conditions [16–18]. Our study revealed some differences in the origin of temperature sensitivity among the TDF mutants. The *rrd1* mutation causes a truncation of the C-terminal domain of the RRD1 protein (Fig. 2A). This finding, together with the enhancement of poly(A)^+^ mitochondrial mRNA accumulation at elevated temperatures (Fig. 3D), implies that, in *rrd1*, RRD1 is partially functional at least at the permissive temperature, and that its activity is more severely affected at the non-permissive temperature. In contrast, the *rrd2* and *rid4-1* mutations introduce a stop codon close to the N-terminus of RRD2 and RID4, respectively, likely resulting in the total loss of their functions (Fig. 2A). The complete abolishment of RNA editing of the RRD2 and RID4 target sites in the *rrd2* and *rid4-1* mutants, regardless of temperature (Fig. 5A and fig. S8A), further supported this idea. However, in *rrd2*, deficient cyt c biogenesis was observed only at high temperature (Fig. 5, C and D). This might be accounted for by the temperature sensitivity of the function of either *ccb2* or *ccb3*, which exhibit alteration of the amino acid sequence in *rrd2*, because of impaired RNA editing (Fig. 5A). In *rid4-1*, a huge reduction in complex V biosynthesis was observed both at permissive and non-permissive temperatures (Fig. 5B). These results suggest that complex V deficiency is more deleterious at higher temperatures, which can explain the temperature sensitivity of the LR fasciation phenotype of *rid4-1*.

### Impaired mitochondrial electron transport causes LR fasciation likely via ROS production

The phenocopy of the LR fasciation phenotype of the TDF mutants by treatment with respiratory inhibitors demonstrated the causal relationship between defective mitochondrial electron transport and excessive cell division during early LR development (Fig. 6, A to D). Mitochondrial electron transport is best known for its role in driving ATP synthesis through oxidative phosphorylation. Given the lack of LR fasciation after treatment with the mitochondrial uncoupler CCCP (Fig. 6, E and F), reduced ATP production seems unlikely to be the cause of LR fasciation. The fact that the huge reduction in complex V levels observed in *rid4* (Fig. 5B) does not lead to LR fasciation at the permissive temperature [18] is also supportive of this idea. Experiments using the ROS inducer PQ and the antioxidant ascorbate (Fig. 6, G and H) pointed to mitochondrial ROS generation as the potential trigger of LR fasciation. A previous study also observed enhanced cell division after the application of another ROS inducer, alloxan, during auxin-induced LR formation [36]. In agreement with this ‘ROS hypothesis’, all three respiratory inhibitors used in our study (rotenone, antimycin A, and oligomycin) are potent inducers of oxidative stress [37].

ROS have been implicated in stress-induced morphogenic responses (SIMR) [38]. Several studies have shown the involvement of phytohormonal regulation in ROS-triggered SIMR. Altered auxin levels and/or distribution have been proposed as potential mediators in the modulation of cell proliferation in response to oxidative stress [36,38]. Several recent studies have found antagonistic interactions between auxin signaling and mitochondrial ROS [39]. Auxin is a critical factor in LR development, and the centripetal auxin-gradient formation in early-stage LR primordia is thought to contribute to the organization of the LR primordium [6,7]. However, neither the pattern nor the intensity of the auxin response visualized by the *DR5::GUS* reporter seemed to be altered under PQ treatment, in contrast to the diffuse pattern observed after the application the auxin polar transport inhibitor NPA (Fig. 6I). This indicates that ROS-induced LR fasciation is not attributable to a failure in auxin-gradient formation. Further studies of LR fasciation caused by oxidative stress will elucidate novel aspects of the control of cell proliferation during plant organogenesis.

### Mitochondrial RNA processing is linked to the control of cell proliferation

Mutants of nuclearly encoded mitochondrial RNA processing factors have proven to be useful in probing the physiological roles of mitochondrial gene expression. In particular, studies of C-to-U editing PPR protein genes have led to a collection of about 100 mutants, among which RNA-editing mutants are available for most mitochondrial genes [35]. The majority of the mutations confer visible phenotypes, such as growth retardation, impaired embryo development, late flowering, or reduced pollen sterility [35]. Similar developmental defects are also observed in mutants of genes encoding other mitochondrial proteins, including *ndufs4* (complex I mutant), *rpoTmp* (RNA polymerase mutant), and *atphb3* (prohibitin mutant) [40]. These results suggest that mitochondria play a supportive role in plant growth, presumably by supplying energy through oxidative phosphorylation. In this study, however, we found that mitochondrial RNA processing is required for preventing excessive cell division during LR primordium formation. This suggests that mitochondrial gene expression not only supports active cell proliferation for growth and development but also participates in the local fine-tuning of organ morphogenesis by restricting cell proliferation.

In summary, our study identified an unexpected link between mitochondrial RNA processing and the control of cell proliferation at the early stage of LR development, probably mediated by changes in the level of mitochondrial ROS. This finding provides a novel clue for the physiological significance of mitochondrial activities in the restrictive regulation of cell proliferation required for the proper morphogenesis of plant organs.

## Materials and Methods

### Plant materials and growth condition

*Arabidopsis thaliana* (L.) Heynh. ecotypes Columbia (Col) and Landsberg *erecta* (L*er*) were used as Arabidopsis in this work. The TDF mutants *rrd1, rrd2*, and *rid4-1* were described previously [16–18]. The *ags1* mutant (*ags1-1*) was also described previously [23]. The *35S::Mt-GFP* line was a gift from Shin-ichi Arimura [41]. *rid4-2* was derived from an ethyl methanesulfonate-mutagenized population of the L*er* strain of Arabidopsis. SALK_027874 was obtained from the Arabidopsis Biological Resource Center. *rrd1* mutant strains harboring either *ags1* or *AGS1*^*c*^ were obtained by *rrd1* (L*er* background) × *ags1* (Col background) and *rrd1* × Col crosses, respectively. The *DR5::GUS* line [42] was a gift from Tom J. Guilfoyle and was crossed three times to L*er* before use. Primers for the genotyping the mutants are listed in table S1.

For tissue culture experiments, donor plants were aseptically grown on Murashige– Skoog medium supplemented with 1.0% (w/v) sucrose, buffered to pH 5.7 with 0.05% (w/v) 2-morpholinoethanesulfonic acid (MES), and solidified with 1.5% (w/v) agar under continuous light (10–15 µmol m^−2^ s^−1^) at 22°C. For observation of seedling phenotypes, plants were aseptically grown on the same medium solidified with 1.5% (w/v) agar or 0.8% (w/v) gellan gum under continuous light (50–80 µmol m^−2^ s^−1^) at 22°C or 28°C. For self-propagation and crossing, plants were grown on vermiculite under continuous light (approximately 50 µmol m^−2^ s^−1^) at 22°C unless otherwise indicated.

### LR induction

As described previously [19], explants were prepared from 4-day-old seedlings grown on agar plates, and cultured on root-inducing medium (RIM) under continuous light (15–25 µmol m^−2^ s^−1^) for the induction of semi-synchronous formation of LRs. RIM was B5 medium supplemented with 2.0% (w/v) glucose and 0.5 mg l^−1^ indole-3-butyric acid, buffered to pH 5.7 with 0.05% (w/v) MES, and solidified with 0.25% (w/v) gellan gum. Culture temperature was set to 22°C for the permissive condition and to 28°C for the non-permissive condition.

### Histological analysis

For whole-mount observation, tissue samples were fixed in 25 mM sodium phosphate buffer (pH 7.0) containing 2% (w/v) formaldehyde and 1% (w/v) glutaraldehyde, rinsed with 100 mM sodium phosphate buffer (pH 7.0), and cleared with an 8:1:2 (w/v/v) mixture of chloral hydrate, glycerin, and water. Observations were made with a microscope equipped with Nomarski optics (BX50-DIC; Olympus) to obtain differential interference contrast (DIC) images.

For morphometric analysis of LR primordia, in order to highlight cell organization, the method of [43] was instead employed for tissue fixation and clearing. Developmental stages of LR primordia were determined according to [43]. LR primordia at Stages II to early III and at Stages IV to V were chosen from samples that had been collected after 16 to 24 hours and 24 to 48 hours of culture in the semi-synchronous root induction system, respectively, and were measured for their width and cell number.

For histochemical detection of GUS reporter expression, tissue samples were fixed in 90% (v/v) acetone overnight at −20°C, rinsed with 100 mM sodium phosphate (pH 7.0), and incubated in X-Gluc solution [0.5 mg ml^−1^ 5-bromo-4-chloro-3-indolyl β-D-glucuronide cyclohexylammonium salt, 0.5 mM potassium ferricyanide, 0.5 mM potassium ferrocyanide, 100 mM sodium phosphate (pH 7.4)] for 140 min at 37 °C. After rinsing with 100 mM sodium phosphate buffer (pH 7.0), the samples were mounted on glass slides with an 8:1:2 (w/v/v) mixture of chloral hydrate, glycerin, and water, and then subjected to DIC microscopy.

### Chromosome mapping

The TDF mutants in the L*er* background were crossed with the wild-type Col strain, and the resultant F_1_ plants were self-pollinated to produce F_2_ seeds or test-crossed with the mutant plants to produce TC_1_ seeds. The TC_2_ lines were then developed by separately collecting self-pollinated progenies from each individual TC_1_ plant. F_2_ plant or TC_2_ lines were checked for the ability of AR formation at 28°C and for DNA polymorphism between L*er* and Col. Chromosome locations of the TDF mutations were determined on the basis of linkage between the mutations and the L*er* alleles of polymorphic marker loci.

### Identification of the TDF genes

Sequencing of the genomic regions to which the TDF mutations were mapped led to identification of candidates of *RRD1, RRD2*, and *RID4* as At3g25430, At1g32415, and At2g33680, respectively. Identification of these genes was confirmed by the complementation test or the allelism test as described below.

For the complementation test, genomic clones GL07, encompassing At3g25430 (2.9-kbp 5’-flanking sequence, 2.6-kbp coding sequence, and 2.5-kbp 3’-flanking sequence), and GL91321, encompassing At2g33680 (1.8-kbp 5’-flanking sequence, 3.5-kbp coding sequence, and 2.0-kbp 3’-flanking sequence), were isolated from a transformation-competent genome library [19], and introduced into the *rrd1* and *rid4* mutants, respectively. The resultant transformants were examined for the ability of AR formation at 28°C. To determine allelism between *rrd2* and SALK_027874, which carries a T-DNA insertion in At1g32415, F_1_ progeny derived by crossing *rrd2* with SALK_027874 was examined for the ability of AR formation at 28°C.

### Plasmid construction

Genomic DNA from L*er* was used as a template for PCR-based amplification of DNA fragments of interest. *RRD1::RRD1:GFP* was constructed by inserting the –2780/+2495 region of the *RRD1* gene (+1 = the first base of the translation initiation codon), which encompassed the genomic region from the promoter to the end of the protein-coding sequence, and the coding sequence of sGFP into pGreen0029 (John Innes Centre). *RID4::RID4:GFP* was similarly constructed by inserting the –2297/+2181 region of the *RID4* gene and the sGFP-coding sequence into pGreen0029. For the construction of *35S::RRD2:GFP*, the +1/+2283 region of the *RRD2* gene was inserted into the pSHO1 vector, a derivative of pHTS13 [44]. Plasmids for the PARN activity assay were constructed by inserting the coding sequence of RRD1 or human PARN (hPARN) into the pHAT vector (Clontech). The hPARN sequence was derived from the GNP Human cDNA clone IRAK071M01 (RIKEN BioResource Research Center). In this plasmid construction, the N-terminal mitochondrial localization signal (24 a.a.) sequence was deleted from the RRD1 coding sequence, and the SEP-tag C9D sequence [45] was added to the C-terminus of both RRD1 and hPARN sequences to improve the solubility of these protein products.

### Plant transformation

DNAs such as reporter gene constructs and genomic fragments were transformed into *Agrobacterium tumefaciens* and then into Arabidopsis by the floral dip method [46] or its modified version [47]. Transgenic plants were selected by antibiotic resistance and genotyped by PCR for the introduction of the correct transgene. Transient expression of *35S::RRD2:GFP* in protoplasts of cultured cells were done as described in [44].

### Expression and localization analysis of GFP reporters

Expression patterns of *RRD1* and *RID4* were examined with transgenic plants harboring *RRD1::RRD1:GFP* and *RID4::RID4:GFP*, respectively. Roots of 6-day-old seedlings of these plants were counterstained with 10 mg l^−1^ of propidium iodide and fluorescence images were obtained using a confocal microscope (FV3000; Olympus). Expression analysis of *35S::Mt-GFP* was performed in the same conditions using a different confocal microscope (FV1200; Olympus). To investigate subcellular localization of the RRD1 and RID4 proteins, protoplasts were prepared from calli that had been induced from the *RRD1::RRD1:GFP* and *RID4::RID4:GFP* explants. The protoplasts were incubated with 100 nM Mitotracker Orange (Invitrogen) for 15 minutes to visualize mitochondria and then observed using the LSM710 system (Carl Zeiss).

### Microarray analysis and data processing

For microarray analysis, total RNA was extracted with TRIzol reagent (Invitrogen) from explants that had been cultured on RIM for 12 hours in the semi-synchronous LR induction system and purified using the RNeasy microkit (QIAGEN). Affymetrix ATH1 microarrays were hybridized with biotinylated cRNA targets prepared from the RNA samples according to the manufacturer’s instructions. It should be noted here that all the targets were derived from poly(A)^+^ RNA in principal because the T7-oligo(dT)_24_ primer was used for reverse-transcription at the first step of target preparation. Experiments were performed in biological triplicates. The data sets obtained were processed with a variant of MAS5.0 utilizing robust radius-minimax estimators [48]. Differential gene expression was identified by RankProd 2.0 [49]. The details of the microarray data was deposited in the Gene Expression Omnibus (http://www.ncbi.nlm.nih.gov/geo/) under accession number GSE34595.

### Analysis of mRNA polyadenylation status with RACE-PAT

RACE-PAT was performed principally according to [28]. Total RNA was extracted with TRIzol reagent (Invitrogen) either from LR-induced explants or seedlings. Total RNA was treated with RNase-free DNase I (Promega) to eliminate genomic DNA, and reverse-transcribed with T7-oligo(dT)_24_ as a primer using the PrimeScript II 1st strand cDNA Synthesis kit (TaKaRa). Then the poly(A) tail status was analyzed by PCR with a combination of gene-specific and T7 promoter primers. The thermal cycling program consisted of initial 2-minute denaturation at 95°C followed by 30 cycles of 20 seconds at 95°C, 20 seconds at 57°C, and 10 seconds at 72°C. Primers for the RACE-PAT are listed in table S1.

### qRT-PCR analysis

For qRT-PCR, total RNA was extracted with TRIzol reagent (Invitrogen) from explants LR-induced at 28°C for 12 hours. To eliminate genomic DNA, total RNA was treated with RNase-free DNase I (Promega), and reverse-transcribed with a random hexamer or oligo(dT)_24_ primer using SYBR Premix ExTaq II (TaKaRa). qRT-PCR reactions were performed with gene-specific forward and reverse primers using the PrimeScript RT-PCR kit (TaKaRa) on the StepOne Real-Time PCR system (Applied Biosystems). The thermal cycling program consisted of initial 30-second denaturation at 95°C followed by 40 cycles of 5 seconds at 95°C and 30 seconds at 60°C. At the end of run, melting curves were established for each PCR product to check the specificity of amplification. Expression levels of mRNAs of interest were normalized relative to *TUBULIN4* (At5g44340) expression. DNA fragments amplified from poly(A)^+^ transcripts of several genes including *cob* were sequenced to check the occurrence of mitochondrial editing, which confirmed that they are derived from the mitochondrial genome but not from their copies present in chromosome 2 [50]. Experiments were performed in biological triplicates. Primers for the qRT-PCR analysis are listed in table S1.

### PARN activity assay of recombinant RRD1

The pHAT plasmids in which the RRD1 or hPARN sequence had been inserted were transformed into the Rosetta-gami 2 strain or the M15 strain of *E. coli*. Colonies were grown overnight at 37°C in LB medium containing 100 µg ml^-1^ ampicillin and 25 µg ml^-1^ chloramphenicol for Rosetta-gami 2 and 100 µg ml^-1^ ampicillin and 25 µg ml^-1^ kanamycin for M15. The cultures were diluted (6:100) in the same medium and grown at 37°C for approximately 3 hours to reach OD_600_ of 0.3 to 0.4, and then treated with 0.2 mM isopropyl *β*-D-1-thiogalactopyranoside (IPTG) overnight at 18°C to induce the production of the his-tagged RRD1 and hPARN proteins. After cell lysis, the proteins were purified by TALON Metal Affinity Resin (Clontech) and filtered with Amicon Ultra 0.5ml (30K; Merck Millipore). For the ribonuclease activity assay, the purified proteins (0.125 mg) or RNase If (1.25 U; NEB) were incubated at 25°C for 60 minutes with a fluorescent-labeled RNA substrate (5’-fluorescein isothiocyanate (FITC)-CUUUUAG(A_20_); this sequence was derived from the 3’ extremity of *cox1* mRNA (Fig. 3C)) in 10 µL of reaction medium (1.5 mM MgCl_2_, 100 mM KCl, 0.1 U RNasin Ribonuclease Inhibitor (Promega), 20 mM HEPES-KOH (pH 7.0), 0.2 mM EDTA, 0.25 mM dithiothreitol, 10% (v/v) glycerol, 0.1% BSA) [51]. The reaction was stopped by adding an equal volume of gel loading mix (90% formamide, 0.5% (w/v) EDTA, 0.025% (w/v) bromophenol blue) and heating to 90°C for 3 minutes before cooling on ice. The reaction mixtures were loaded onto a 7 M urea-16% polyacrylamide gel and separated by electrophoresis.

### CR-RT PCR analysis of the 3’ end of mRNA

CR-RT PCR analysis was performed principally according to [27]. Total RNA was extracted with TRIzol reagent (Invitrogen) from seedlings that had been cultured for 7 days at 22°C and then 2 days at 28°C. To eliminate genomic DNA, total RNA was treated with DNase I (RT grade; Nippon Gene). Next 1 µg of total RNA was circularized with T4 RNA ligase (Promega), desalted with Amicon Ultra 0.5ml (10K; Merck Millipore), and then reverse-transcribed with a *cox1* specific primer (Atcox1-1; table S1) using M-MLV (Moloney Murine Leukemia Virus) Reverse Transcriptase (RNase H minus, point mutant; Promega). The RNA template was degraded by adding 1/5 volume of 1 M NaOH to the reaction mixture and incubating at room temperature for 10 minutes. The solution was neutralized by adding 1 M HCl and the cDNA was purified with the illustra GFX PCR DNA and Gel Band Purification Kit (GE Healthcare). The 5’-3’ junction sequence was amplified by PCR with *cox1* specific primers Atcox1-5’(−176..-196) and Atcox1-3’(+17..+38) using Ex Taq Hot Start Version (Takara). The thermal cycling program consisted of initial 4 minute-denaturation at 95°C, followed by 40 cycles of 20 seconds at 95°C, 20 seconds at 50°C, and 40 seconds at 72°C. The PCR products were purified with the Wizard SV Gel and PCR Clean-Up System (Promega) and cloned into the pGEM-T Easy Vector (Promega) using DNA Ligation Kit <Mighty Mix> (Takara). The constructed vector was transformed into the DH5*α* strain of *E*.*coli*, and about 20 clones were sequenced. Primers for the CR RT-PCR analysis are listed in table S1.

### Analysis of mitochondrial mRNA editing

For the analysis of mitochondrial mRNA editing, total RNA was extracted with TRIzol reagent (Invitrogen) from explants LR-induced at 28°C for 12 hours. Total RNA was treated with RNase-free DNase I (Promega), and reverse-transcribed with a random hexamer using the PrimeScript II 1st strand cDNA Synthesis kit (TaKaRa). Gene specific primers were used to amplify cDNA by PCR using Ex Taq Hot Start Version (Takara). The thermal cycling program consisted of initial 4-minute denaturation at 95°C followed by 30 to 40 cycles of 30 seconds at 95°C, 30 seconds at 55°C, and 90 to 120 seconds at 72°C. The PCR products were purified either by ExoStar DNA purification reagent (GE Healthcare) or Wizard SV Gel and PCR Clean-Up System (Promega), and then sequenced.

### Analysis of mitochondrial protein

Isolation of intact mitochondria was performed principally according to [52]. Seed-derived callus cultured in liquid callus-inducing medium (CIM) [16,17] in the dark with gentle shaking was used as starting material. About 16 g of callus was homogenized in 40 ml ice-cold grinding buffer (0.3 M Mannitol, 50 mM Tetrasodium pyrophosphate, 2 mM EDTA (Disodium salt), 0.5 % (w/v) PVP-40, 0.5 % (w/v) BSA, 20 mM L-cysteine, pH 8.0 (HCl)) with a mortar, pestle, and glass beads (0.4-mm diameter). The homogenate was filtered through four layers of Miracloth (Millipore) and centrifuged at 2,300g for 5 minutes twice. The resulting supernatant was centrifuged at 18,000g for 10 minutes. The resulting pellet was resuspended in wash buffer (0.3 M Mannitol, 10 mM *N*-Tris(hydroxymethyl)methyl-2-aminoethanesulfonic acid (TES), 0.1% (w/v) BSA, pH 7.5 (NaOH)) and layered over a three-step Percoll (GE Healthcare) gradient (40%, 21%, and 16% (v/v)). The gradient was centrifuged at 23,500 rpm (approximately 40,000g to 70,000g) for 30 minutes. Mitochondria were collected from the 21% and 40% interface and washed twice in wash buffer (without BSA) by centrifugation at 18,000g for 10 minutes.

For BN-PAGE analysis, 10 µg protein of mitochondria was solubilized in 12 µL Native PAGE Sample Buffer (1% n-dodecyl-β-D-maltoside (DDM), Thermo Fisher Scientific), mixed with 1.8 µL of sample additive (33.3% (w/v) glycerol, 1.67% (w/v) Coomassie Brilliant Blue (CBB) G250), and then separated by electrophoresis on a NativePAGE 4 to 16%, Bis-Tris Gel (Thermo Fisher Scientific).

For immunoblot analysis, proteins separated via SDS–PAGE were transferred to a PVDF membrane and exposed to a primary antibody against cyt c (AS08 343A, Agrisera; 1:5000 dilution). As a secondary antibody, we used a peroxidase-labeled anti-rabbit antibody (NIF824, GE Healthcare; 1:5000 dilution). Immunodetection was performed by incubating the membranes in the Western BLoT Quant HRP Substrate (Takara) and recording the chemiluminescence by LuminoGraphI(ATTO).

### Graph drawing

Bar charts were drawn using KaleidaGraph (Abelbeck Software) or Excel for Mac (Microsoft). Scatter plots were drawn using KaleidaGraph or R software. Dot plots were drawn using R software. Violin plots were overlayed using the geom_flat_violin function (https://gist.github.com/JoachimGoedhart/98ec16c041aab8954083097796c2fe81). Box plots were drawn using R software.

## Supplementary Materials

Fig. S1. Chromosome mapping of the TDF mutations, *rrd1, rrd2*, and *rid4-1*.

Fig. S2. Complementation analysis and allelism test for the identification of the TDF genes RRD1, RRD2, and RID4.

Fig. S3. Identification and characterization of the *rid4-2* mutant.

Fig. S4. Functionality and expression of *RRD1::RRD1:GFP* and *RID4::RID4:GFP*

Fig. S5. Characterization of RRD1 function.

Fig. S6. Sequence alignment of RRD1 and PARNs of various organisms.

Fig. S7. Comprehensive analysis of mitochondrial mRNA editing in *rrd2* and *rid4-1*.

Fig. S8. Analysis of mitochondrial mRNA editing in *rrd2* and *rid4-1*.

Fig. S9. Effects of NPA and PQ on LR formation. Table S1. Primers used in this study.

## Acknowledgments

We thank Tsuyoshi Nakagawa for providing the binary vector pGW3, Mamoru Sugita for valuable discussion on the PPR proteins, Hajime Sakurai for the technical support for the expression of *35S::RRD2:GFP* in protoplasts, Shin-ichi Arimura for providing the *35S::Mt-GFP* line, Yuta Otsuka and Yuki Kondo for the assistance on GFP imaging, Yukiko Sugisawa for the technical support for microarray data collection, Hatsune Morinaka for the assistance on qRT-PCR, and Tom J. Guilfoyle for providing the *DR5::GUS* line, and the RIKEN BioResource Research Center for providing the hPARN cDNA clone.

## Funding

This work was supported by Grant-in-Aid for Scientific Research on Priority Areas (No. 19060001 to M.S.), the Graduate Program for Leaders in Life Innovation (GPLLI) at the University of Tokyo Life Innovation Leading Graduate School (for A.M.) from the Ministry of Education, Culture, Sports, Science and Technology, Japan (MEXT), and by Grant-in-Aid for Scientific Research (B) (No. 25291057 to M.S.) and Grants-in-Aid for JSPS Fellows (No. 09J08676 to K.O. and No. 17J05722 to A.M.) from Japan Society for the Promotion of Science (JSPS).

## Author contributions

K.O. designed and performed experiments and data analysis mostly in the first half of this study, including histological analysis of fasciated LRs, positional cloning of *RRD1* and *RRD2*, construction of the reporter genes, subcellular localization analysis of the TDF proteins, microarray data collection, initial analysis of the poly(A) status of mitochondrial mRNAs, and initial pharmacological analysis with respiratory inhibitors. A.M. designed and performed experiments and data analysis mostly in the latter half of this study, including expression analysis of the TDF genes, microarray data mining, analysis of polyadenylation and editing of mitochondrial mRNAs, genetic analysis with *ags1*, analysis of mitochondrial proteins, and pharmacological analysis with respiration- and ROS-related drugs. M.K. identified *RID4* by positional cloning. M.N. designed and performed analysis of PARN activity of recombinant RRD1. A.K. conducted chromosome mapping of *rrd1* and some of the initial characterization of the TDF phenotype. H.T. isolated the *rid4-2* mutant. M.A. and M.Sa. conducted chromosome mapping of *rid4-2*. K.Y. collected preliminary data on the genetic relationship between *rrd1* and *ags1*. T.Ha. and K.N. contributed to the research design and data interpretation for mitochondrial respiration-related analysis. T.U. contributed to the research design, imaging analysis of GFP reporters, and data interpretation for subcellular localization. Y.Y., T.N., and K.Y. performed preliminary analysis of RNA editing. Y.Y., T.K., and T.N. contributed to the research design and data interpretation for RNA editing-related analysis. Y.S. contributed to the analysis of the recombinant RRD1 protein. T.Hi. contributed to the research design and data interpretation for RNA metabolism-related analysis. M.Su. launched and directed the study and conducted preliminary analyses. K.O., A.M., and M.Su. wrote the manuscript. All authors read and approved the paper.

## Competing interests

The authors declare no competing financial interests.

## Data and materials availability

The microarray data has been deposited in the Gene Expression Omnibus (http://www.ncbi.nlm.nih.gov/geo/) under accession number GSE34595. All data needed to evaluate the conclusions in the paper are present in the paper and/or the Supplementary Materials. Plant materials used in this study can be distributed upon request to the corresponding author.

## Figures and Tables

**Fig. S1 Chromosome mapping of the TDF mutations, *rrd1, rrd2*, and *rid4-1*. (A)** Chromosome mapping of the *rrd1* mutation. The black rectangles represent annotation units around the *RRD1* locus on chromosome 3, and the red numerals correspond to the number of recombination events between DNA polymorphism markers and the *RRD1* locus. The *rrd1* mutation was mapped to the region covered by the annotation units MJL12, MTE24, and MWL2. Sequencing of this region, followed by complementation analysis, identified the *rrd1* mutation as a G-to-A transition in At3g25430 (orange arrow). **(B)** Chromosome mapping of the *rrd2* mutation. The black rectangles represent annotation units around the *RRD2* locus on chromosome 1, and the red numerals correspond to the number of recombination events between DNA polymorphism markers and the *RRD2* locus. The *rrd2* mutation was mapped to the region covered by the annotation units F3C3, F27G20, and F5D14. Sequencing of this region, followed by allelism analysis, identified the *rrd2* mutation as a G-to-A transition in At1g32415 (orange arrow). **(C)** Chromosome mapping of the *rid4-1* mutation. The black rectangles represent the annotation units around the *RID4* locus on chromosome 2, and the red numerals correspond to the number of recombination events between DNA polymorphism markers and the *RID4* locus. The *rid4* mutation was mapped to the region covered by the annotation units F4P9 and T1B8. Sequencing of this region, followed by complementation analysis, identified the *rid4* mutation as a G-to-A transition in At2g33680 (orange arrow).

**Fig. S2 Complementation analysis and allelism test for the identification of the TDF genes *RRD1, RRD2*, and *RID4*. (A)** Complementation analysis for the identification of the *RRD1* gene. The genomic fragment GL07 encompassing At3g25430, where an *rrd1* phenotype-linked mutation was found, was introduced into the wild-type (WT) plant, which was then crossed with *rrd1*. Each individual of the F2 progeny was genotyped for the *rrd1* allele and the GL07 transgene. Hypocotyl explants of the F2 progeny were cultured on RIM at 28°C for 14 days and examined for adventitious rooting. The explants were categorized according to the length of the ARs (Short, shorter than 5 mm; Long, longer than 5 mm) and counted. The results showed that the development of ARs, which was highly temperature sensitive in the *rrd1* mutant, was clearly rescued by the introduction of GL07. Therefore, we concluded that the *RRD1* gene corresponds to At3g25430. **(B and C)** Defect of AR formation in a T-DNA insertion mutant of At1g32415. SALK_027874 carries a T-DNA insertion in the middle of At1g32415 (**B**). The transcribed region and open reading frame of At1g32415 are indicated by the open arrow and grey box, respectively. Hypocotyl explants of Col, L*er, rrd2*, and SALK_027874 were cultured on RIM at 28°C for 27 days and examined for adventitious rooting (**C**). The results indicated that SALK_027874 and *rrd2* are defective in AR formation at this temperature. As AR formation was not significantly affected at 22°C in both *rrd2* and SALK_027874 (data not shown), SALK_027874 was shown to be temperature sensitive for root development, as was *rrd2*. Bar, 1 cm. **(D)** Allelism test for the identification of the *RRD2* gene. The *rrd2* mutant was crossed with SALK_027874 carrying a T-DNA insertion in At1g32415, in which an *rrd2* phenotype-linked mutation was found. Each individual of the F2 progeny was genotyped for the *rrd2* and the T-DNA insertion alleles. Hypocotyl explants of the F2 progeny were cultured on RIM at 28°C for 14 days and examined for adventitious rooting. The explants were categorized according to the length of the ARs (Short, shorter than 5 mm; Long, longer than 5 mm) and counted. The results indicated clearly that *rrd2* and SALK_027874 are allelic. Therefore, we concluded that the *RRD2* gene corresponds to At1g32415. **(E)** Complementation analysis for the identification of the *RID4* gene. The genomic fragment GL91321 encompassing At2g33680, where we found an *rid4* phenotype-linked mutation, was introduced into *rid4*, and the resultant transgenic *rid4* mutant harboring GL91321 (*rid4*/GL91321) was used for complementation analysis. Hypocotyl explants of the WT plant, *rid4*, and *rid4*/GL91321-2 were cultured on RIM at 28°C for 19 days and examined for AR formation. The development of ARs, which was highly temperature sensitive in the *rid4* mutant, was clearly rescued by the introduction of GL91321. Therefore, we concluded that the *RID4* gene corresponds to At2g33680. Scale bar, 1 cm.

**Fig. S3 Identification and characterization of the *rid4-2* mutant. (A)** Representative images of LRs formed at 22°C or 28°C in the explants of the wild-type plant or the *rid4-2* mutant after 6 days of culture. Fasciated LRs were observed in *rid4-2* explants at 28°C. **(B)** Phenotypes of seedlings that were grown for 7 days on vertical agar plates. Seedlings were grown either at 22°C or 28°C. **(C)** Allelism test between *rid4-1* and *rid4-2*. F_1_ plants derived from a reciprocal crossing between *rid4-1* and *rid4-2* were subjected to phenotypic analysis regarding AR formation. Hypocotyl explants of *rid4-1, rid4-2*, L*er* WT, and F_1_ plants were cultured on RIM for 24 days at 28°C. **(D)** Chromosome mapping of the *rid4-2* mutation. The black rectangles represent annotation units around the *RID4* locus on chromosome 2, and the red numerals correspond to the number of recombination events between DNA polymorphism markers and the *RID4* locus. *rid4-2* was mapped to the region covered by the annotation units F4P9 and T29F13. Sequencing of this region, followed by complementation analysis, identified the *rid4-2* mutation as a G-to-A transition in At2g33680 (orange arrow). Scale bars, 100 μm (**A**) and 1 cm (**B**).

**Fig. S4 Functionality and expression of *RRD1::RRD1:GFP* and *RID4::RID4:GFP*. (A)** Phenotypic complementation of *rrd1* by the introduction of the GFP reporter gene *RRD1::RRD1:GFP*. F3 plants homozygous for *rrd1* derived from the cross between rrd1 and Col carrying *RRD1::RRD1:GFP* were phenotyped for AR formation from hypocotyl explants at 28°C and genotyped for the presence of *RRD1::RRD1:GFP* (+, present; –, absent). **(B)** Phenotypic complementation of *rid4* by the introduction of the GFP reporter gene *RID4::RID4:GFP. rid4* homozygotes in which the genetic background had been partially replaced by crossing with the Col strain were transformed with *RID4::RID4:GFP*. Plants of the resultant T2 line were phenotyped for AR formation from hypocotyl explants at 28°C and genotyped for the presence of *RID4::RID4:GFP* (+, present; –, absent). **(C)** Expression patterns of *RRD1* and *RID4* in the root apical region. GFP signals in the primary roots of the transgenic plants harboring *RRD1::RRD1:GFP* and *RID4::RID4:GFP* revealed strong expression of *RRD1* and *RID4* in the root apical meristem. Scale bar, 100 μm.

**Fig. S5 Characterization of RRD1 function. (A–C)** Microarray analysis of mitochondrial genes in *rrd1*. MA plot for the microarray analysis of poly(A)^+^ transcripts of *rrd1* vs. wild-type (WT) explants in which LRs were induced at 28°C for 12 hours. The characterized mitochondrial genes are shown in blue, while the noncharacterized mitochondrial ORFs or pseudogenes are shown in red (**A**). The names of the characterized genes are indicated in (**B**). **(C)** The function and array element ID of the characterized mitochondrial genes in the GeneChip Arabidopsis Genome ATH1 Array. **(D and E)** PARN activity assay of RRD1. TALON-purified fraction of the total cell lysate with or without IPTG induction (**D**). PARN activity assay using the TALON-purified fraction **(E)**.

**Fig. S6 Sequence alignment between RRD1 and PARNs from various organisms**. An alignment of amino acid sequences was generated between RRD1 and PARNs from humans, *Xenopus laevis*, and Arabidopsis using the ClustalW program and processed with BOXSHADE (http://www.ch.embnet.org/software/BOX_form.html). Identical and similar amino acid residues are highlighted on black and grey backgrounds, respectively. The R3H domain is marked by the dotted orange box, and the RNA recognition motif (RRM) is marked by the solid blue box. These domains are conserved in the animal PARNs (human and *Xenopus laevis*), but are not clearly present in the Arabidopsis PARN and RRD1. The pink asterisk represents the tryptophan codon that was changed to a stop codon by the *rrd1* mutation. The red arrowheads represent the four residues that are important for PARN activity (*29*).

**Fig. S7 Comprehensive analysis of mitochondrial mRNA editing in *rrd2* and *rid4-1***. A sequencing analysis of mitochondrial mRNA editing was performed using explants in which LRs were induced at 28°C for 12 hours. The color code indicates the level of C-to- U RNA editing at each site (editing status: –0.5 = 100% C, 0.5 = 100% U). The presumptive specific editing sites of RRD2 and RID4 are marked by solid black boxes. RNA editing in *rid4-2* was also analyzed for sites affected in *rid4-1*.

**Fig. S8 Analysis of mitochondrial mRNA editing in *rrd2* and *rid4-1*. (A)** A sequencing analysis of mitochondrial mRNA editing was performed using explants in which LRs were induced at 22°C for 12 hours. **(B)** Alignment of the estimated binding sequence of RRD2 and RID4 (*31*). **(C)** Analysis of the mRNA editing of *ccb3* in explants that were cultured at 22°C or 28°C for 12 hours.

**Fig. S9 Effects of NPA and PQ on LR formation**. Root explants 6 days after LR induction under treatment with NPA or PQ. Scale bar, 1 mm.

**Table S1. Primers used in this study**.

## Notes

https://www.ncbi.nlm.nih.gov/geo/query/acc.cgi?acc=GSE34595

### Competing Interest Statement

The authors have declared no competing interest.

## References and Notes

1. Torres-Martínez, H.H., Rodríguez-Alonso, G., Shishkova, S., and Dubrovsky, J.G. (2019). Lateral root primordium morphogenesis in angiosperms. Front. Plant Sci. 10, 206.

2. Hammani, K., and Giegé, P. (2014). RNA metabolism in plant mitochondria. Trends Plant Sci. 19, 380–389.

3. von Wangenheim, D., Fangerau, J., Schmitz, A., Smith, R.S., Leitte, H., Stelzer, E.H.K., and Maizel, A. (2016). Rules and self-organizing properties of post-embryonic plant organ cell division patterns. Curr. Biol. 26, 439–449.

4. Goh, T., Toyokura, K., Wells, D.M., Swarup, K., Yamamoto, M., Mimura, T., Weijers, D., Fukaki, H., Laplaze, L., Bennett, M.J., et al. (2016). Quiescent center initiation in the *Arabidopsis* lateral root primordia is dependent on the *SCARECROW* transcription factor. Development 143, 3363–3371.

5. Lavenus, J., Goh, T., Roberts, I., Guyomarc’h, S., Lucas, M., De Smet, I., Fukaki, H., Beeckman, T., Bennett, M., and Laplaze, L. (2013). Lateral root development in *Arabidopsis*: fifty shades of auxin. Trends Plant Sci. 18, 455–463.

6. Benková, E., Michniewicz, M., Sauer, M., Teichmann, T., Seifertová, D., Jürgens, G., and Friml, J. (2003). Local, efflux-dependent auxin gradients as a common module for plant organ formation. Cell 115, 591–602.

7. Geldner, N., Richter, S., Vieten, A., Marquardt, S., Torres-Ruiz, R.A., Mayer, U., and Jürgens, G. (2004). Partial loss-of-function alleles reveal a role for GNOM in auxin transport-related, post-embryonic development of *Arabidopsis*. Development 131, 389–400.

8. De Smet, I., Lau, S., Voß, U., Vanneste, S., Benjamins, R., Rademacher, E.H., Schlereth, A., De Rybel, B., Vassileva, V., Grunewald, W., et al. (2010). Bimodular auxin response controls organogenesis in *Arabidopsis*. Proc. Natl. Acad. Sci. U. S. A. 107, 2705–2710.

9. De Smet, I., Vassileva, V., De Rybel, B., Levesque, M.P., Grunewald, W., Van Damme, D., Van Noorden, G., Naudts, M., Van Isterdael, G., De Clercq, R., et al. (2008). Receptor-like kinase ACR4 restricts formative cell divisions in the *Arabidopsis* root. Science. 322, 594–597.

10. Murphy, E., Vu, L.D., den Broeck, L., Lin, Z.F., Ramakrishna, P., van de Cotte, B., Gaudinier, A., Goh, T., Slane, D., Beeckman, T., et al. (2016). RALFL34 regulates formative cell divisions in *Arabidopsis* pericycle during lateral root initiation. J. Exp. Bot. 67, 4863–4875.

11. Hirota, A., Kato, T., Fukaki, H., Aida, M., and Tasaka, M. (2007). The auxin-regulated AP2/EREBP gene PUCHI is required for morphogenesis in the early lateral root primordium of *Arabidopsis*. Plant Cell 19, 2156–2168.

12. Du, Y.J., and Scheres, B. (2017). PLETHORA transcription factors orchestrate de novo organ patterning during *Arabidopsis* lateral root outgrowth. Proc. Natl. Acad. Sci. U. S. A. 114, 11709–11714.

13. Benitez-Alfonso, Y., Faulkner, C., Pendle, A., Miyashima, S., Helariutta, Y., and Maule, A. (2013). Symplastic intercellular connectivity regulates lateral root patterning. Dev. Cell 26, 136–147.

14. Napsucialy-Mendivil, S., Alvarez-Venegas, R., Shishkova, S., and Dubrovsky, J.G. (2014). *ARABIDOPSIS HOMOLOG of TRITHORAX1 (ATX1)* is required for cell production, patterning, and morphogenesis in root development. J. Exp. Bot. 65, 6373–6384.

15. Vermeer, J.E.M., von Wangenheim, D., Barberon, M., Lee, Y., Stelzer, E.H.K., Maizel, A., and Geldner, N. (2014). A spatial accommodation by neighboring cells is required for organ initiation in *Arabidopsis*. Science. 343, 178–183.

16. Sugiyama, M. (2003). Isolation and initial characterization of temperature-sensitive mutants of *Arabidopsis thaliana* that are impaired in root redifferentiation. Plant Cell Physiol. 44, 588–596.

17. Konishi, M., and Sugiyama, M. (2003). Genetic analysis of adventitious root formation with a novel series of temperature-sensitive mutants of *Arabidopsis thaliana*. Development 130, 5637–5647.

18. Otsuka, K., and Sugiyama, M. (2012). Tissue organization of fasciated lateral roots of Arabidopsis mutants suggestive of the robust nature of outer layer patterning. J. Plant Res. 125, 547–554.

19. Ohtani, M., Demura, T., and Sugiyama, M. (2010). Particular significance of SRD2-dependent snRNA accumulation in polarized pattern generation during lateral root development of Arabidopsis. Plant Cell Physiol. 51, 2002–2012.

20. Hu, Z.B., Vanderhaeghen, R., Cools, T., Wang, Y., De Clercq, I., Leroux, O., Nguyen, L., Belt, K., Millar, A.H., Audenaert, D., et al. (2016). Mitochondrial defects confer tolerance against cellulose deficiency. Plant Cell 28, 2276–2290.

21. Pavlopoulou, A., Vlachakis, D., Balatsos, N.A.A., and Kossida, S. (2013). A comprehensive phylogenetic analysis of deadenylases. Evol. Bioinforma. 9, 491–497.

22. Lee, D., Park, D., Park, J.H., Kim, J.H., and Shin, C. (2019). Poly(A)-specific ribonuclease sculpts the 3 ‘ends of microRNAs. RNA 25, 388–405.

23. Hirayama, T., Matsuura, T., Ushiyama, S., Narusaka, M., Kurihara, Y., Yasuda, M., Ohtani, M., Seki, M., Demura, T., Nakashita, H., et al. (2013). A poly(A)-specific ribonuclease directly regulates the poly(A) status of mitochondrial mRNA in *Arabidopsis*. Nat. Commun. 4.

24. Kanazawa, M., Ikeda, Y., Nishihama, R., Yamaoka, S., Lee, N.H., Yamato, K.T., Kohchi, T., and Hirayama, T. (2020). Regulation of the poly(A) status of mitochondrial mRNA by poly(A)-specific ribonuclease is conserved among land plants. Plant Cell Physiol. 61, 470–480.

25. Hirayama, T. (2014). A unique system for regulating mitochondrial mRNA poly(A) status and stability in plants. Plant Signal. Behav. 9, 1–4.

26. Holec, S., Lange, H., Canaday, J., and Gagliardi, D. (2008). Coping with cryptic and defective transcripts in plant mitochondria. Biochim. Biophys. Acta 1779, 566–573.

27. Forner, J., Weber, B., Thuss, S., Wildum, S., and Binder, S. (2007). Mapping of mitochondrial mRNA termini in *Arabidopsis thaliana*: t-elements contribute to 5 ‘and 3 ‘end formation. Nucleic Acids Res. 35, 3676–3692.

28. Sallés, F.J., Richards, W.G., and Strickland, S. (1999). Assaying the polyadenylation state of mRNAs. Methods-a Companion to Methods Enzymol. 17, 38–45.

29. Reverdatto, S. V., Dutko, J.A., Chekanova, J.A., Hamilton, D.A., and Belostotsky, D.A. (2004). mRNA deadenylation by PARN is essential for embryogenesis in higher plants. RNA 10, 1200–1214.

30. Barkan, A., and Small, I. (2014). Pentatricopeptide Repeat Proteins in Plants. Annu. Rev. Plant Biol. 65, 415–442.

31. Kobayashi, T., Yagi, Y., and Nakamura, T. (2019). Comprehensive prediction of target RNA editing sites for PLS-class PPR proteins in *Arabidopsis thaliana*. Plant Cell Physiol. 60, 862–874.

32. Cheng, S.F., Gutmann, B., Zhong, X., Ye, Y.T., Fisher, M.F., Bai, F.Q., Castleden, I., Song, Y., Song, B., Huang, J.Y., et al. (2016). Redefining the structural motifs that determine RNA binding and RNA editing by pentatricopeptide repeat proteins in land plants. Plant J. 85, 532–547.

33. Giegé, P., Grienenberger, J.M., and Bonnard, G. (2008). Cytochrome c biogenesis in mitochondria. Mitochondrion 8, 61–73.

34. Heazlewood, J.L., Whelan, J., and Millar, A.H. (2003). The products of the mitochondrial orf25 and orfB genes are Fo components in the plant F1Fo ATP synthase. Febs Lett. 540, 201–205.

35. Takenaka, M., Jörg, A., Burger, M., and Haag, S. (2019). RNA editing mutants as surrogates for mitochondrial SNP mutants. Plant Physiol. Biochem. 135, 310–321. Available at: https://doi.org/10.1016/j.plaphy.2018.12.014.

36. Pasternak, T., Potters, G., Caubergs, R., and Jansen, M.A.K. (2005). Complementary interactions between oxidative stress and auxins control plant growth responses at plant, organ, and cellular level. J. Exp. Bot. 56, 1991–2001.

37. Willems, P., Mhamdi, A., Stael, S., Storme, V., Kerchev, P., Noctor, G., Gevaert, K., and Van Breusegem, F. (2016). The ROS wheel: refining ROS transcriptional footprints. Plant Physiol. 171, 1720–1733.

38. Potters, G., Pasternak, T.P., Guisez, Y., and Jansen, M.A.K. (2009). Different stresses, similar morphogenic responses: integrating a plethora of pathways. Plant Cell Environ. 32, 158–169.

39. Huang, S.B., Van Aken, O., Schwarzländer, M., Belt, K., and Millar, A.H. (2016). The roles of mitochondrial reactive oxygen species in cellular signaling and stress response in plants. Plant Physiol. 171, 1551–1559.

40. Van Aken, O., Whelan, J., and Van Breusegem, F. (2010). Prohibitins: mitochondrial partners in development and stress response. Trends Plant Sci. 15, 275–282. Available at: http://dx.doi.org/10.1016/j.tplants.2010.02.002.

41. Arimura, S., and Tsutsumi, N. (2002). A dynamin-like protein (ADL2b), rather than FtsZ, is involved in *Arabidopsis* mitochondrial division. Proc. Natl. Acad. Sci. U. S. A. 99, 5727–5731.

42. Ulmasov, T., Murfett, J., Hagen, G., and Guilfoyle, T.J. (1997). Aux/IAA Proteins repress expression of reporter genes containing natural and highly active synthetic auxin response elements. Plant Cell 9, 1963–1971.

43. Malamy, J.E., and Benfey, P.N. (1997). Organization and cell differentiation in lateral roots of *Arabidopsis thaliana*. Development 124, 33–44.

44. Ueda, T., Yamaguchi, M., Uchimiya, H., and Nakano, A. (2001). Ara6, a plant-unique novel type Rab GTPase, functions in the endocytic pathway of *Arabidopsis thaliana*. EMBO J. 20, 4730–4741.

45. Kato, A., Maki, K., Ebina, T., Kuwajima, K., Soda, K., and Kuroda, Y. (2007). Mutational analysis of protein solubility enhancement using short peptide tags. Biopolymers 85, 12–18.

46. Clough, S.J., and Bent, A.F. (1998). Floral dip: a simplified method for Agrobacterium-mediated transformation of *Arabidopsis thaliana*. Plant J. 16, 735–743.

47. Martinez-Trujillo, M., Limones-Briones, V., Cabrera-Ponce, J.L., and Herrera-Estrella, L. (2004). Improving transformation efficiency of *Arabidopsis thaliana* by modifying the floral dip method. Plant Mol. Biol. Report. 22, 63–70.

48. Kohl, M., and Deigner, H.P. (2010). Preprocessing of gene expression data by optimally robust estimators. BMC Bioinformatics 11.

49. Del Carratore, F., Jankevics, A., Eisinga, R., Heskes, T., Hong, F., and Breitling, R. (2017). RankProd 2.0: a refactored bioconductor package for detecting differentially expressed features in molecular profiling datasets. Bioinformatics 33.

50. Stupar, R.M., Lilly, J.W., Town, C.D., Cheng, Z., Kaul, S., Buell, C.R., and Jiang, J.M. (2001). Complex mtDNA constitutes an approximate 620-kb insertion on *Arabidopsis thaliana* chromosome 2: implication of potential sequencing errors caused by large-unit repeats. Proc. Natl. Acad. Sci. U. S. A. 98, 5099–5103.

51. Cheng, Y., Liu, W.-F., Yan, Y.-B., and Zhou, H.-M. (2005). A nonradioactive assay for poly(A)-specific ribonuclease activity by methylene blue colorimetry. Protein Pept. Lett. 13, 125–128.

52. Murcha, M.W., and Whelan, J. (2015). Isolation of intact mitochondria from the model plant species Arabidopsis thaliana and Oryza sativa. In Methods in Molecular Biology (Humana Press Inc.), pp. 1–12. Available at: http://link.springer.com/10.1007/978-1-4939-2639-8_1.

