## Supplemental material for "Temperature-dependent fasciation mutants connect mitochondrial RNA processing to control of lateral root morphogenesis"

### Supplementary figure 1

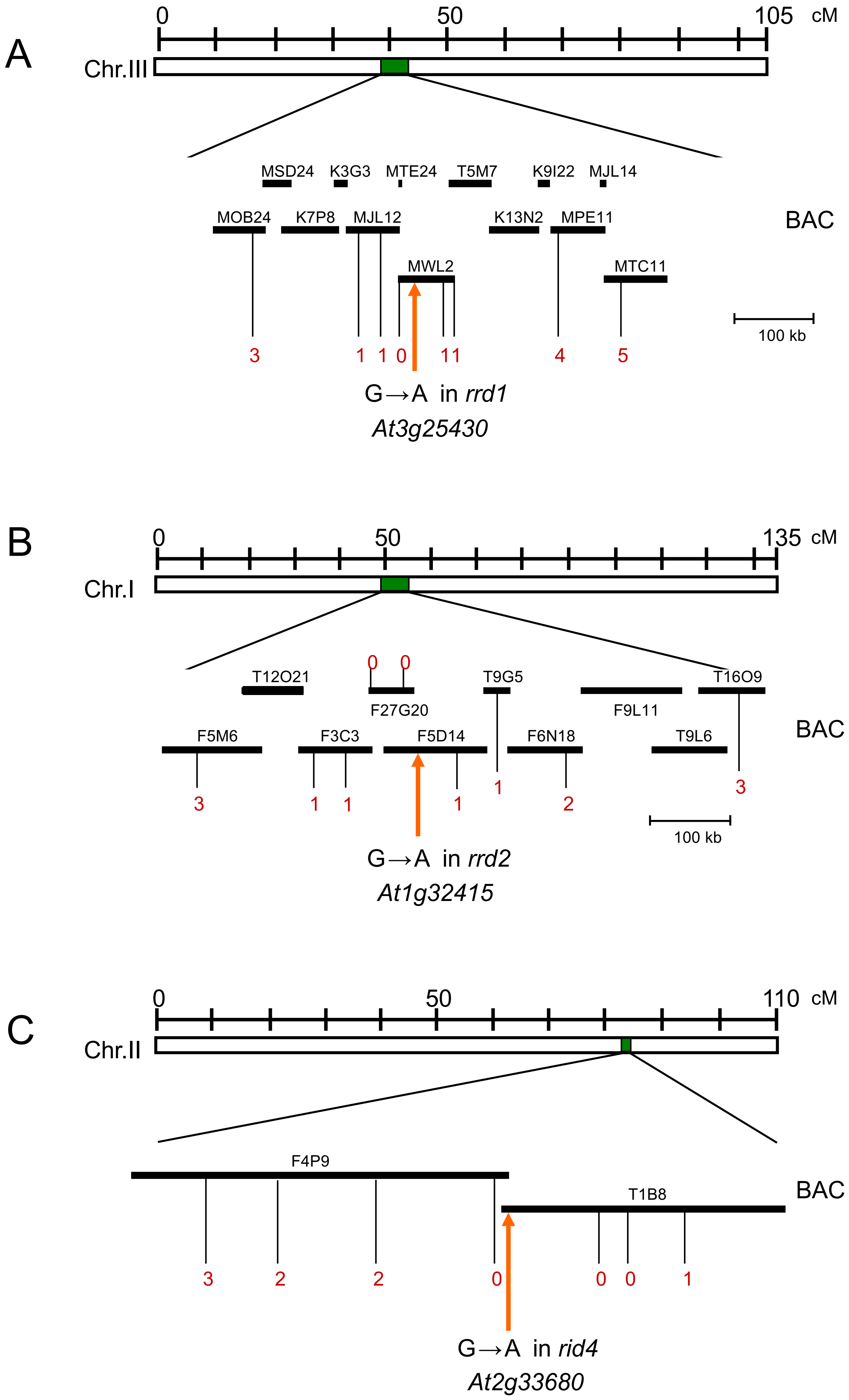

### Supplementary figure 2

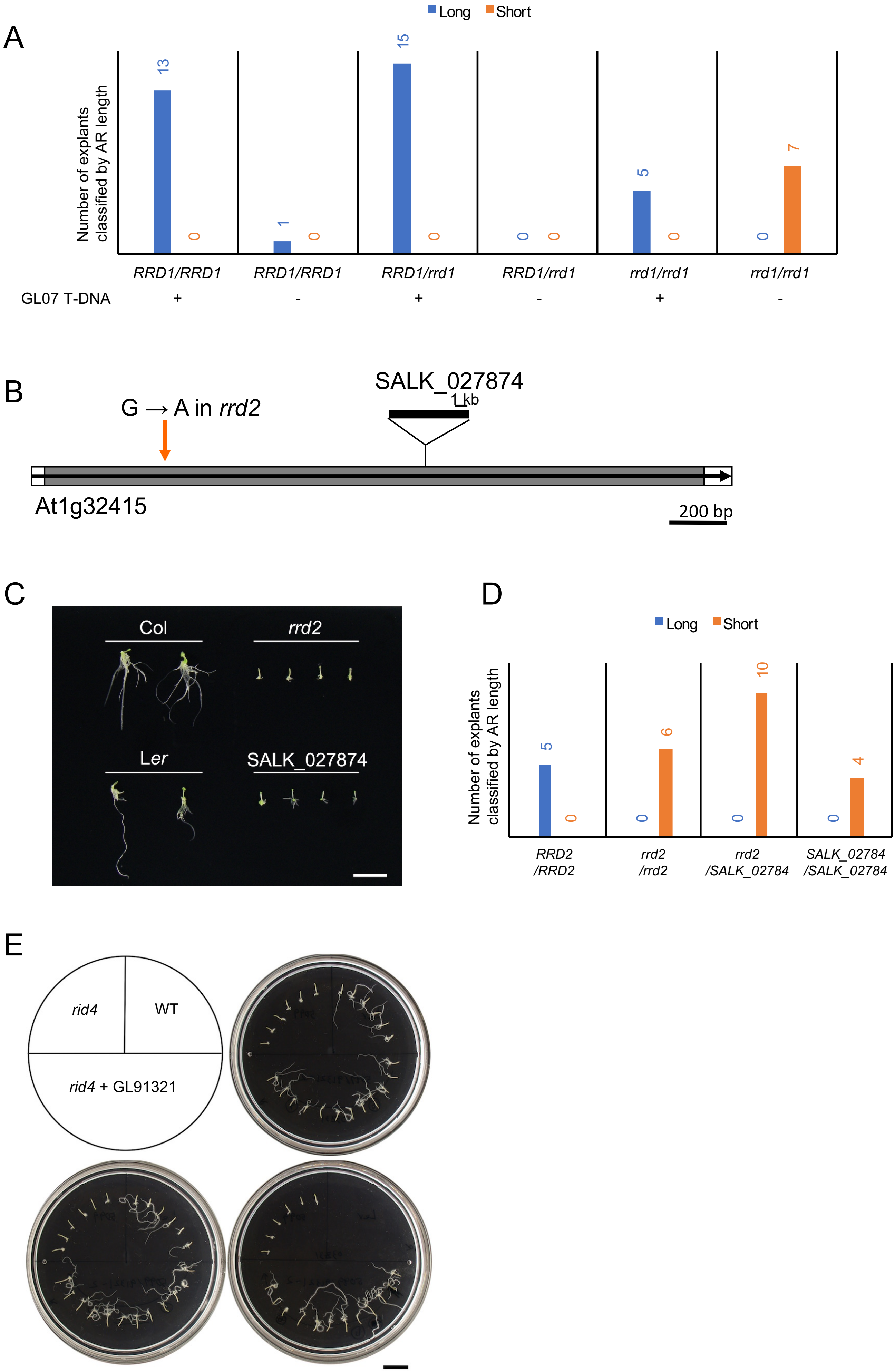

**Supplementary figure 3**

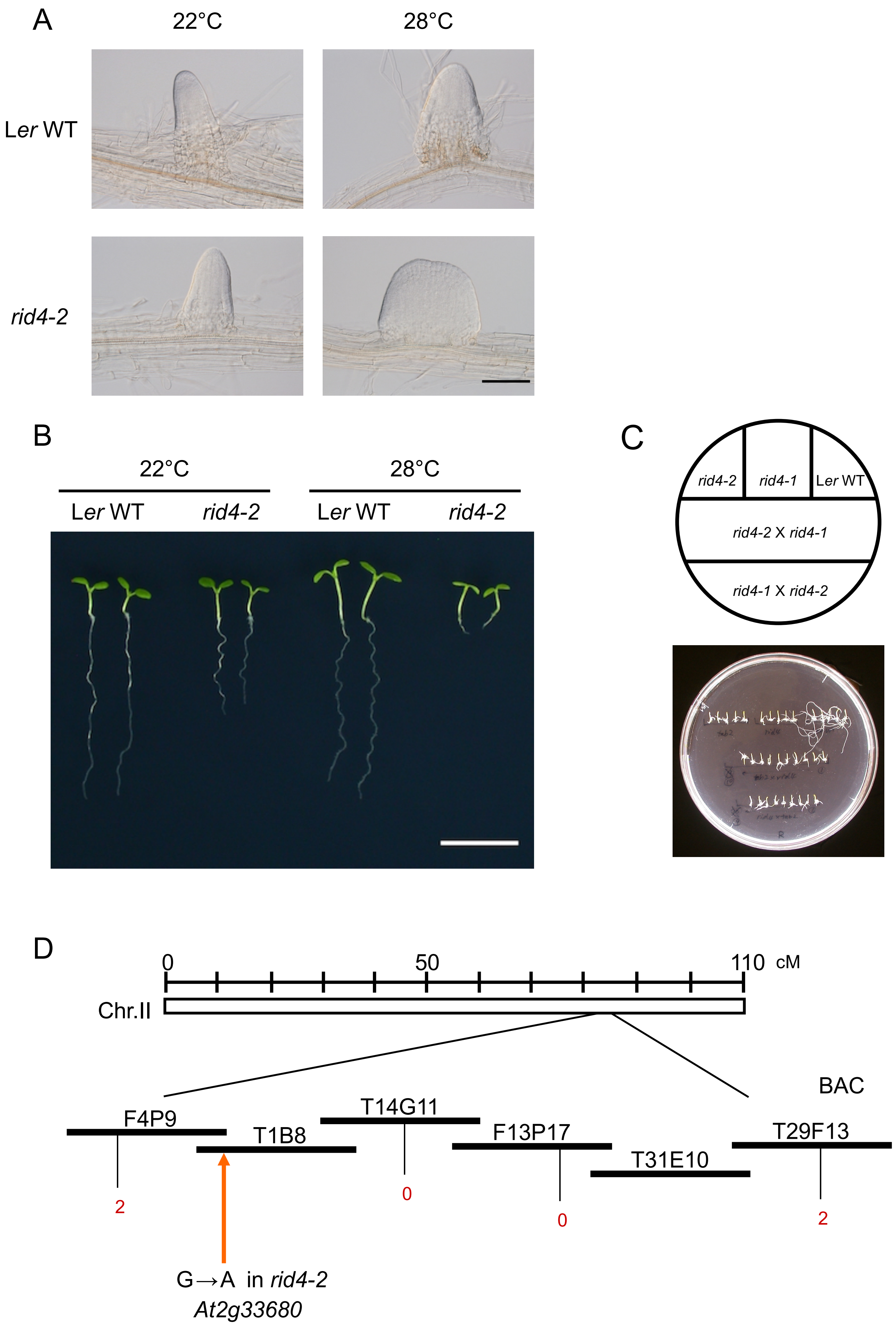

### Supplementary figure 4

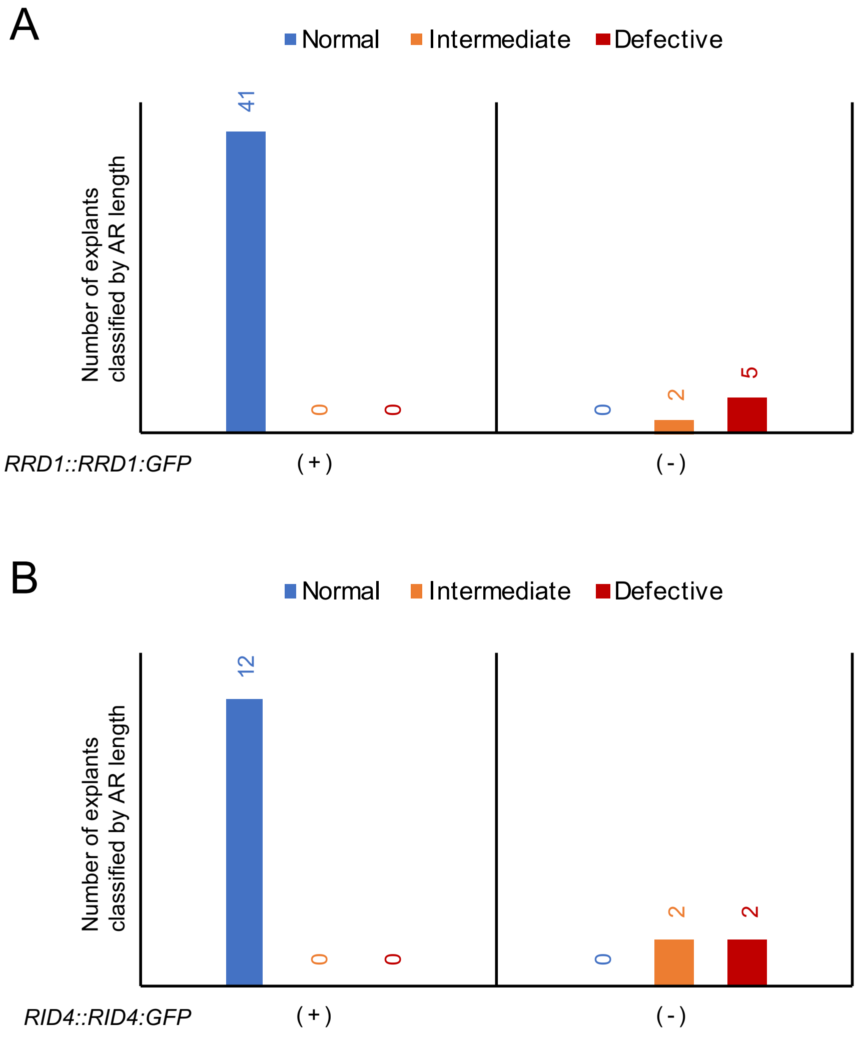


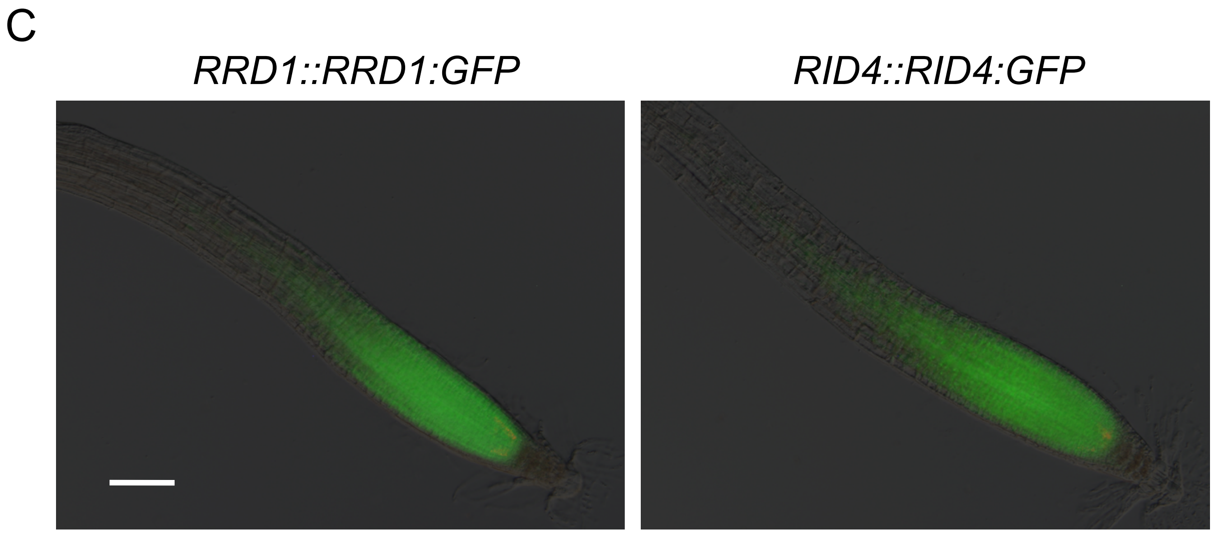

### Supplementary figure 5

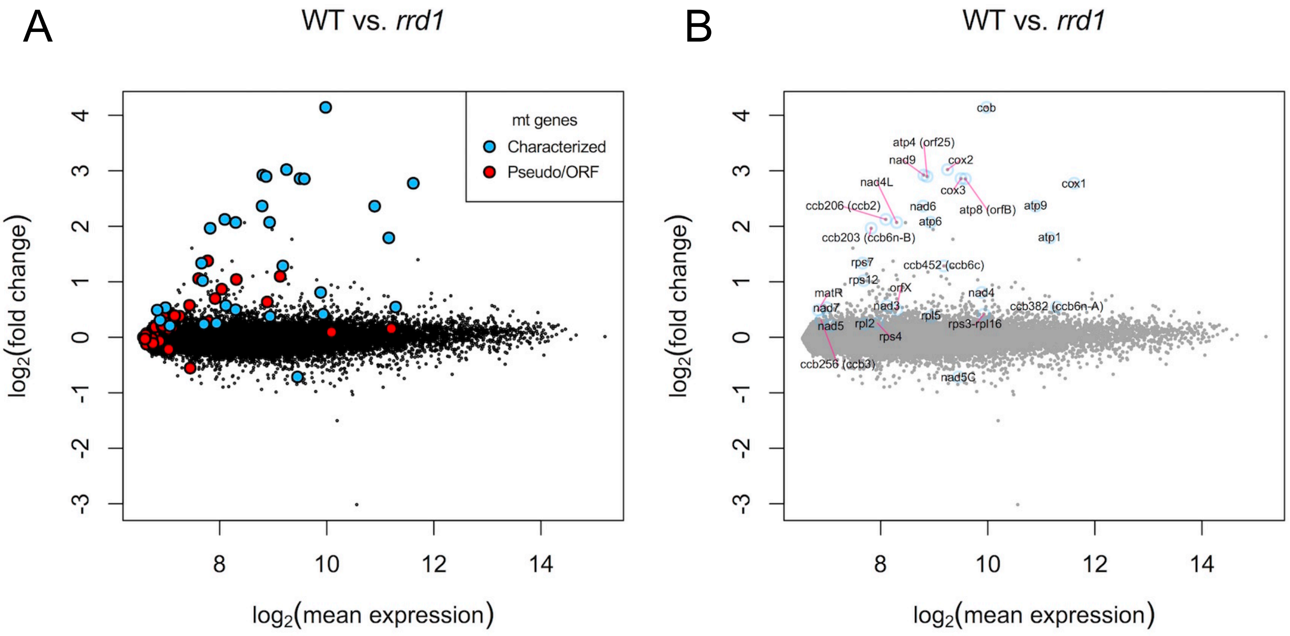


C

| Gene | Function | ID |
|---|---|---|
| nad3 | complex I | 244920_s_at |
| nad4 | complex I | 244929_at |
| nad4L | complex I | 244902_at |
| nad5 | complex I | 257337_at |
| nad5C | complex I | 257338_s_at |
| nad6 | complex I | 244953_s_at |
| nad7 | complex I | 244925_at |
| nad9 | complex I | 244943_at |
| cob | complex III | 266045_s_at |
| cox1 | complex IV | 257333_at |

| Gene | Function | ID |
|---|---|---|
| cox2 | complex IV | 244950_at |
| cox3 | complex IV | 263509_s_at |
| atp1 | complex V | 265228_s_at |
| atp4 (orf25) | complex V | 244901_at |
| atp6 | complex V | 266012_s_at |
| atp8 (orfB) | complex V | 265230_s_at |
| atp9 | complex V | 257339_s_at |
| ccb203 (ccb6n-B) | cytC biogenesis | 244919_at |
| ccb206 (ccb2) | cytC biogenesis | 244945_at |
| ccb256 (ccb3) | cytC biogenesis | 263510_s_at |

| Gene | Function | ID |
|---|---|---|
| ccb382 (ccb6n-A) | cytC biogenesis | 244912_at |
| ccb452 (ccb6c) | cytC biogenesis | 244951_s_at |
| rpl2 | Ribosome | 265232_s_at |
| rpl5 | Ribosome | 266044_s_at |
| rps3-rpl16 | Ribosome | 244944_s_at |
| rps4 | Ribosome | 266042_s_at |
| rps7 | Ribosome | 265238_s_at |
| rps12 | Ribosome | 263502_s_at |
| matR | Maturase | 244926_s_at |
| orfX | Protein transport ? | 244928_s_at |

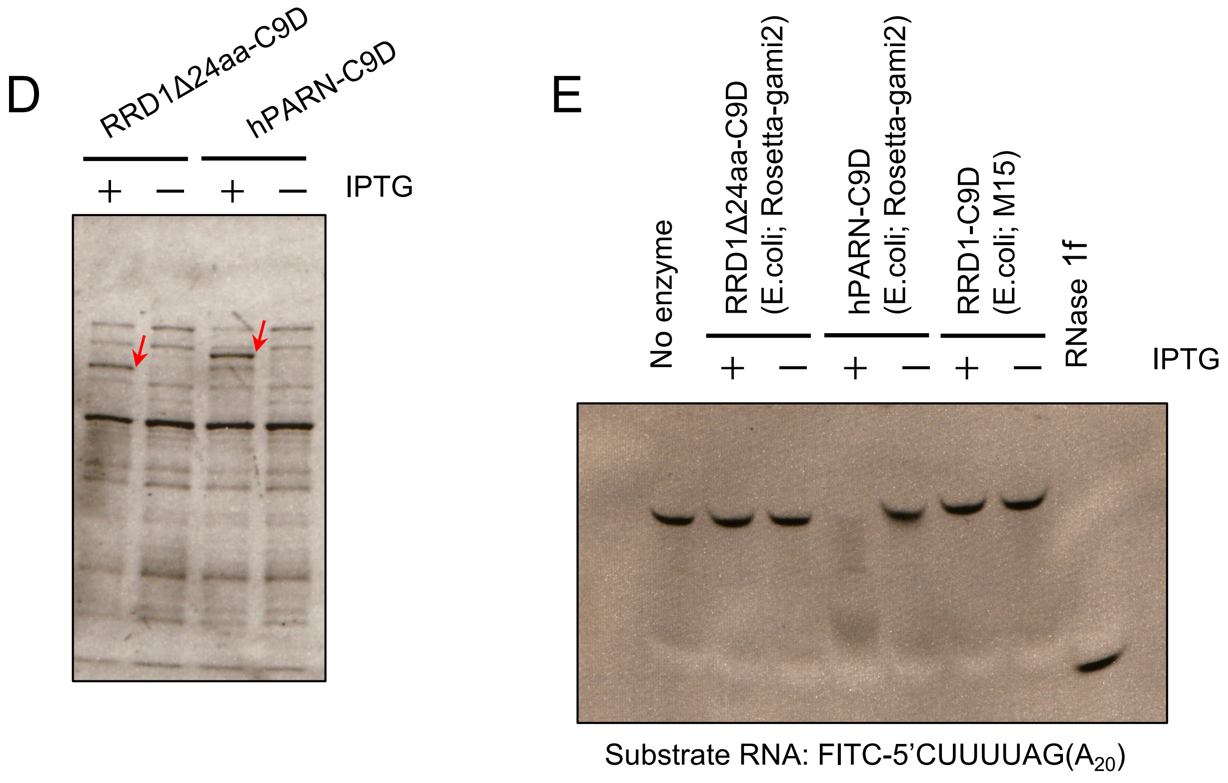

### Supplementary figure 6

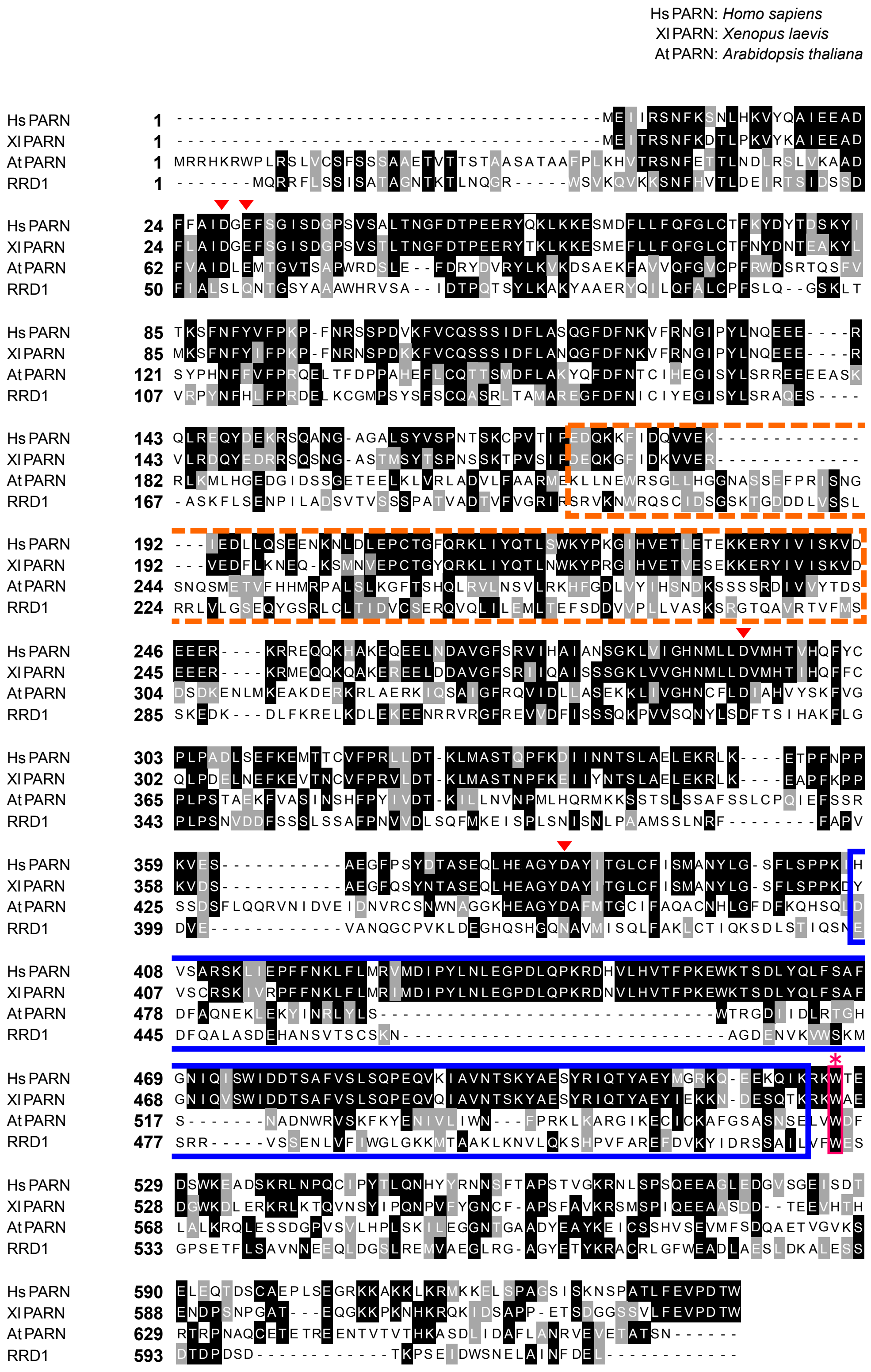

### Supplementary figure 7 (1 of 3)

| *atp1* | 1110 | 1178 | 1292 | 1415 | 1484 |
|---|---|---|---|---|---|
| LerWT | -0.36 | 0.49 | 0.49 | 0.49 | 0.50 |
| *rrd2* | -0.40 | 0.49 | 0.49 | 0.46 | 0.50 |
| *rid4-1* | -0.42 | 0.48 | 0.47 | 0.46 | 0.50 |

| *atp4* | 89 | 138 | 215 | 248 | 250 | 251 | 382 | 395 | 416 |
|---|---|---|---|---|---|---|---|---|---|
| LerWT | -0.23 | 0.31 | 0.50 | 0.45 | 0.10 | 0.48 | -0.40 | 0.50 | 0.47 |
| *rrd2* | -0.24 | 0.32 | 0.50 | 0.47 | 0.12 | 0.49 | -0.45 | 0.48 | 0.46 |
| *rid4-1* | -0.31 | 0.25 | 0.48 | 0.46 | 0.04 | 0.47 | -0.42 | -0.48 | 0.45 |
| *rid4-2* | -0.31 | 0.22 | 0.48 | 0.45 | -0.04 | 0.48 | -0.41 | -0.28 | 0.47 |

| *atp6* | 475 |
|---|---|
| LerWT | 0.46 |
| *rrd2* | 0.43 |
| *rid4* | 0.42 |

| *atp9* | 53 | 83 | 167 | 224 |
|---|---|---|---|---|
| LerWT | 0.46 | 0.49 | 0.48 | 0.46 |
| *rrd2* | 0.46 | 0.47 | 0.48 | 0.46 |
| *rid4-1* | 0.44 | 0.47 | 0.45 | 0.45 |

| *ccb2* | 16 | 28 | 71 | 75 | 78 | 80 | 128 | 137 | 148 | 149 | 154 | 159 | 160 | 164 | 172 | 179 | 181 | 188 | 193 | 194 |
|---|---|---|---|---|---|---|---|---|---|---|---|---|---|---|---|---|---|---|---|---|
| LerWT | -0.04 | 0.09 | 0.01 | -0.48 | -0.49 | -0.06 | 0.03 | 0.04 | 0.04 | 0.01 | 0.00 | 0.04 | 0.10 | 0.02 | 0.12 | 0.01 | 0.01 | -0.48 | -0.02 | 0.00 |
| *rrd2* | -0.19 | -0.22 | -0.45 | -0.50 | -0.50 | -0.29 | -0.22 | -0.23 | -0.25 | -0.26 | -0.26 | -0.24 | -0.24 | -0.22 | -0.21 | -0.21 | -0.22 | -0.48 | -0.25 | -0.31 |
| *rid4-1* | -0.28 | -0.36 | -0.31 | -0.50 | -0.50 | -0.38 | -0.28 | -0.33 | -0.36 | -0.36 | -0.37 | -0.36 | -0.35 | -0.35 | -0.33 | -0.36 | -0.35 | -0.48 | -0.37 | -0.41 |

| *ccb2* | 286 | 304 | 338 | 367 | 379 | 380 | 406 | 424 | 428 | 467 | 475 | 476 | 485 | 512 | 514 | 551 | 554 | 566 | 569 |
|---|---|---|---|---|---|---|---|---|---|---|---|---|---|---|---|---|---|---|---|
| LerWT | 0.03 | 0.03 | 0.01 | 0.00 | -0.04 | 0.02 | -0.42 | -0.01 | -0.02 | -0.06 | 0.02 | 0.01 | 0.02 | -0.05 | 0.00 | 0.01 | 0.00 | -0.02 | 0.03 |
| *rrd2* | -0.19 | -0.24 | -0.20 | -0.23 | -0.25 | -0.21 | -0.45 | -0.26 | -0.23 | -0.27 | -0.21 | -0.22 | -0.21 | -0.29 | -0.24 | -0.23 | -0.27 | -0.25 | -0.21 |
| *rid4-1* | -0.39 | -0.38 | -0.35 | -0.39 | -0.40 | -0.37 | -0.47 | -0.40 | -0.39 | -0.40 | -0.39 | -0.39 | -0.38 | -0.45 | -0.38 | -0.38 | -0.41 | -0.40 | -0.42 |

| *ccb3* | 103 | 133 | 179 | 184 | 262 | 331 | 395 | 400 | 421 | 436 | 446 | 458 | 463 | 467 | 473 | 477 | 497 | 521 | 548 | 568 |
|---|---|---|---|---|---|---|---|---|---|---|---|---|---|---|---|---|---|---|---|---|
| LerWT | 0.26 | 0.22 | 0.22 | 0.23 | -0.46 | 0.27 | 0.25 | 0.21 | 0.25 | 0.24 | 0.28 | 0.28 | 0.27 | 0.28 | 0.31 | -0.48 | 0.34 | 0.33 | 0.35 | 0.32 |
| *rrd2* | -0.37 | -0.18 | -0.25 | -0.37 | -0.50 | -0.23 | -0.28 | -0.23 | -0.28 | -0.21 | -0.24 | -0.22 | -0.27 | -0.26 | -0.27 | -0.45 | -0.24 | -0.27 | -0.38 | -0.24 |
| *rid4-1* | -0.39 | -0.28 | -0.26 | -0.40 | -0.47 | -0.27 | -0.31 | -0.33 | -0.26 | -0.28 | -0.25 | -0.28 | -0.30 | -0.28 | -0.27 | -0.50 | -0.30 | -0.26 | -0.28 | -0.27 |

| *ccb3* | 575 | 608 | 614 | 618 | 619 | 624 | 650 | 656 | 673 |
|---|---|---|---|---|---|---|---|---|---|
| LerWT | 0.36 | 0.37 | 0.34 | 0.40 | 0.37 | 0.02 | 0.38 | 0.39 | 0.20 |
| *rrd2* | -0.43 | -0.31 | -0.30 | -0.36 | -0.31 | -0.45 | -0.41 | -0.33 | -0.40 |
| *rid4-1* | -0.28 | -0.31 | -0.30 | -0.39 | -0.32 | -0.50 | -0.39 | -0.31 | -0.43 |

| *ccb6c* | 50 | 103 | 122 | 123 | 146 | 155 | 160 | 175 | 333 | 334 | 378 | 406 | 415 | 561 | 925 | 1150 | 1172 | 1215 | 1246 | 1280 | 1327 |
|---|---|---|---|---|---|---|---|---|---|---|---|---|---|---|---|---|---|---|---|---|---|
| LerWT | 0.44 | 0.46 | 0.48 | 0.46 | 0.46 | 0.49 | 0.39 | 0.29 | 0.42 | 0.50 | -0.19 | 0.48 | 0.48 | -0.44 | -0.35 | -0.48 | 0.42 | -0.03 | 0.37 | 0.45 | 0.31 |
| *rrd2* | 0.47 | 0.48 | 0.50 | 0.43 | 0.47 | 0.50 | 0.41 | 0.23 | 0.38 | 0.50 | 0.01 | 0.50 | 0.39 | -0.33 | -0.50 | -0.47 | 0.47 | -0.06 | 0.28 | 0.48 | 0.48 |
| *rid4-1* | 0.47 | 0.49 | 0.50 | 0.28 | 0.48 | 0.48 | 0.43 | 0.21 | 0.27 | 0.50 | -0.14 | 0.50 | 0.39 | -0.37 | -0.43 | -0.45 | 0.47 | -0.08 | 0.34 | 0.42 | 0.50 |

| *ccb6n-A* | 44 | 104 | 143 | 157 | 200 | 262 | 269 | 289 | 340 | 371 | 378 | 404 | 484 | 579 | 581 | 709 | 710 | 719 | 779 | 791 | 806 | 955 |
|---|---|---|---|---|---|---|---|---|---|---|---|---|---|---|---|---|---|---|---|---|---|---|
| LerWT | 0.37 | 0.48 | 0.39 | 0.38 | -0.48 | 0.48 | 0.43 | 0.41 | -0.47 | -0.44 | 0.37 | -0.40 | -0.45 | -0.48 | -0.50 | 0.01 | 0.42 | -0.48 | 0.47 | 0.42 | 0.45 | 0.47 |
| *rrd2* | 0.35 | 0.47 | 0.35 | 0.36 | -0.48 | 0.47 | 0.42 | 0.39 | -0.47 | -0.43 | 0.35 | -0.38 | -0.46 | -0.50 | -0.48 | -0.03 | 0.39 | -0.48 | 0.46 | 0.39 | 0.45 | 0.45 |
| *rid4-1* | 0.30 | 0.46 | 0.38 | 0.31 | -0.48 | 0.40 | 0.42 | 0.37 | -0.47 | -0.45 | 0.35 | -0.38 | -0.45 | -0.50 | -0.48 | -0.16 | 0.37 | -0.49 | 0.47 | 0.41 | 0.43 | 0.47 |

| *ccb6n-B* | 65 | 176 | 208 | 226 | 259 | 277 | 301 | 320 | 344 | 356 | 391 | 467 |
|---|---|---|---|---|---|---|---|---|---|---|---|---|
| LerWT | 0.41 | 0.40 | 0.37 | 0.36 | 0.32 | 0.34 | -0.49 | 0.09 | 0.30 | 0.29 | 0.35 | -0.48 |
| *rrd2* | 0.29 | 0.37 | 0.27 | 0.28 | 0.22 | 0.24 | -0.49 | 0.03 | 0.23 | 0.19 | 0.29 | -0.50 |
| *rid4-1* | 0.23 | 0.34 | 0.21 | 0.20 | 0.16 | 0.19 | -0.48 | -0.01 | 0.15 | 0.12 | 0.23 | -0.50 |

| *cob* | 118 | 286 | 325 | 568 | 610 | 853 | 908 | 924 | 982 | 1084 |
|---|---|---|---|---|---|---|---|---|---|---|
| LerWT | 0.48 | 0.48 | 0.43 | 0.45 | -0.28 | 0.50 | 0.50 | -0.38 | 0.50 | 0.50 |
| *rrd2* | 0.48 | 0.49 | 0.49 | 0.48 | -0.26 | 0.41 | 0.50 | -0.37 | 0.41 | 0.46 |
| *rid4-1* | 0.46 | 0.48 | 0.42 | 0.46 | -0.26 | 0.42 | 0.41 | -0.33 | 0.38 | 0.48 |

| *cox2* | 24 | 25 | 27 | 71 | 138 | 253 | 261 | 278 | 379 | 476 | 557 | 581 | 698 | 721 | 742 |
|---|---|---|---|---|---|---|---|---|---|---|---|---|---|---|---|
| LerWT | -0.43 | -0.04 | 0.40 | 0.46 | -0.02 | 0.49 | -0.46 | 0.50 | 0.50 | 0.50 | 0.48 | 0.39 | 0.50 | 0.47 | -0.44 |
| *rrd2* | -0.39 | -0.05 | 0.42 | 0.49 | 0.03 | 0.50 | -0.45 | 0.50 | 0.47 | 0.50 | 0.50 | 0.41 | 0.50 | 0.45 | -0.50 |
| *rid4-1* | -0.39 | -0.02 | 0.38 | 0.46 | 0.10 | 0.50 | -0.45 | 0.50 | 0.50 | 0.50 | 0.50 | 0.45 | 0.50 | 0.49 | -0.49 |

| *cox3* | 112 | 245 | 257 | 311 | 314 | 413 | 422 | 603 |
|---|---|---|---|---|---|---|---|---|
| LerWT | 0.50 | 0.48 | 0.50 | 0.50 | 0.48 | 0.50 | 0.46 | -0.50 |
| *rrd2* | 0.50 | 0.50 | 0.47 | 0.50 | 0.50 | 0.50 | 0.50 | -0.50 |
| *rid4-1* | 0.50 | 0.50 | 0.46 | 0.46 | 0.44 | 0.30 | 0.29 | -0.28 |

Editing status = U / (C+U) - 0.5

| | | | | | | | | | | |
|---|---|---|---|---|---|---|---|---|---|---|
| -0.5 | -0.4 | -0.3 | -0.2 | -0.1 | 0 | 0.1 | 0.2 | 0.3 | 0.4 | 0.5 |

### Supplementary figure 7 (2 of 3)

| *mat R* | 80 | 241 | 284 | 374 | 461 | 1593 | 1596 | 1676 | 1730 | 1731 | 1751 | 1771 | 1807 | 1895 |
|---|---|---|---|---|---|---|---|---|---|---|---|---|---|---|
| LerWT | -0.46 | -0.46 | -0.49 | 0.48 | 0.47 | -0.21 | -0.32 | -0.47 | 0.36 | 0.12 | 0.35 | 0.31 | 0.21 | 0.41 |
| *rrd2* | -0.48 | -0.46 | -0.49 | 0.43 | 0.44 | -0.19 | -0.33 | -0.47 | 0.36 | 0.14 | 0.34 | 0.27 | 0.19 | 0.40 |
| *rid4-1* | -0.48 | -0.46 | -0.48 | 0.42 | 0.44 | -0.29 | -0.39 | -0.46 | 0.32 | 0.04 | 0.30 | 0.28 | 0.15 | 0.38 |

| *nad1a* | 2 | 167 | 265 | 307 | 308 | 376 |
|---|---|---|---|---|---|---|
| LerWT | 0.47 | 0.50 | 0.47 | 0.50 | 0.50 | 0.48 |
| *rrd2* | 0.49 | 0.45 | 0.45 | 0.45 | 0.45 | 0.48 |
| *rid4-1* | 0.42 | 0.43 | 0.45 | 0.45 | 0.44 | 0.48 |

| *nad1b* | 490 | 492 | 493 | 500 | 536 | 546 | 571 | 580 | 635 |
|---|---|---|---|---|---|---|---|---|---|
| LerWT | 0.50 | 0.45 | 0.44 | 0.50 | 0.50 | -0.50 | 0.47 | 0.50 | 0.50 |
| *rrd2* | 0.50 | 0.50 | 0.50 | 0.34 | 0.50 | -0.50 | 0.44 | 0.46 | 0.31 |
| *rid4-1* | 0.50 | 0.50 | 0.50 | 0.23 | 0.50 | -0.50 | 0.45 | 0.43 | 0.41 |

| *nad1c* | 674 | 725 | 743 | 755 | 763 | 797 | 823 | 898 | 928 | 937 |
|---|---|---|---|---|---|---|---|---|---|---|
| LerWT | 0.48 | 0.47 | 0.50 | 0.50 | -0.50 | -0.50 | 0.50 | 0.50 | 0.48 | 0.47 |
| *rrd2* | 0.49 | 0.46 | 0.50 | 0.50 | -0.50 | -0.50 | 0.50 | 0.50 | 0.50 | 0.48 |
| *rid4-1* | 0.48 | 0.50 | 0.50 | 0.50 | -0.50 | -0.50 | 0.50 | 0.50 | 0.48 | 0.47 |

| *nad2a* | 28 | 59 | 89 | 90 | 285 | 341 | 344 | 389 | 394 | 400 | 427 | 441 | 461 | 528 | 530 | 558 |
|---|---|---|---|---|---|---|---|---|---|---|---|---|---|---|---|---|
| LerWT | 0.42 | 0.48 | 0.45 | 0.26 | -0.40 | 0.50 | 0.50 | 0.44 | 0.46 | 0.50 | 0.50 | -0.50 | 0.44 | -0.50 | 0.50 | -0.50 |
| *rrd2* | 0.44 | 0.48 | 0.48 | 0.23 | -0.35 | 0.50 | 0.47 | 0.44 | 0.47 | 0.50 | 0.50 | -0.50 | 0.47 | -0.50 | 0.50 | -0.40 |
| *rid4-1* | 0.45 | 0.48 | 0.46 | 0.24 | -0.34 | 0.47 | 0.50 | 0.41 | 0.47 | 0.46 | 0.50 | -0.42 | 0.47 | -0.38 | 0.50 | -0.45 |

| *nad2b* | 642 | 695 | 821 | 842 | 953 | 961 | 991 | 995 | 1091 | 1160 | 1233 | 1279 | 1280 | 1309 | 1433 | 1436 | 1490 |
|---|---|---|---|---|---|---|---|---|---|---|---|---|---|---|---|---|---|
| LerWT | -0.46 | 0.50 | 0.50 | 0.50 | 0.50 | 0.50 | 0.50 | 0.50 | 0.50 | 0.50 | -0.42 | 0.45 | 0.46 | 0.50 | 0.50 | 0.50 | -0.42 |
| *rrd2* | -0.44 | 0.50 | 0.50 | 0.50 | 0.50 | 0.50 | 0.50 | 0.50 | 0.50 | 0.50 | -0.39 | 0.50 | 0.50 | 0.50 | 0.50 | 0.50 | -0.45 |
| *rid4-1* | -0.44 | 0.50 | 0.50 | 0.50 | 0.50 | 0.50 | 0.50 | 0.50 | 0.50 | 0.50 | -0.43 | 0.50 | 0.50 | 0.50 | 0.50 | 0.50 | -0.40 |

| *nad3* | 8 | 26 | 64 | 69 | 83 | 149 | 211 | 212 | 250 | 254 | 347 | 352 |
|---|---|---|---|---|---|---|---|---|---|---|---|---|
| LerWT | -0.44 | 0.46 | 0.40 | -0.49 | 0.35 | 0.36 | 0.31 | 0.38 | 0.29 | 0.30 | -0.46 | -0.46 |
| *rrd2* | -0.36 | 0.45 | 0.27 | -0.46 | 0.30 | 0.32 | 0.25 | 0.31 | 0.21 | 0.26 | -0.46 | -0.43 |
| *rid4-1* | -0.25 | 0.40 | 0.19 | -0.48 | 0.21 | 0.24 | 0.15 | 0.22 | 0.11 | 0.13 | -0.47 | -0.30 |

| *nad4* | 29 | 74 | 84 | 107 | 124 | 158 | 164 | 166 | 197 | 317 | 362 | 376 | 402 | 403 | 418 | 436 | 437 | 449 | 608 | 659 |
|---|---|---|---|---|---|---|---|---|---|---|---|---|---|---|---|---|---|---|---|---|
| LerWT | 0.45 | 0.43 | -0.45 | 0.45 | 0.48 | 0.44 | 0.46 | 0.43 | 0.47 | 0.49 | 0.44 | 0.50 | -0.50 | 0.50 | -0.50 | 0.50 | 0.50 | 0.50 | 0.45 | 0.50 |
| *rrd2* | 0.45 | 0.36 | -0.33 | 0.50 | 0.50 | 0.50 | 0.45 | 0.46 | 0.50 | 0.50 | 0.50 | 0.50 | -0.50 | 0.50 | -0.50 | 0.50 | 0.50 | 0.50 | 0.50 | 0.50 |
| *rid4-1* | 0.50 | 0.43 | -0.30 | 0.50 | 0.50 | 0.50 | 0.47 | 0.47 | 0.47 | 0.50 | 0.50 | 0.50 | -0.50 | 0.31 | -0.50 | 0.50 | 0.50 | 0.50 | 0.50 | 0.40 |

| | 767 | 784 | 836 | 896 | 977 | 1010 | 1033 | 1101 | 1129 | 1131 | 1133 | 1148 | 1172 | 1206 | 1355 | 1373 | 1401 | 1405 | 1417 | 1433 |
|---|---|---|---|---|---|---|---|---|---|---|---|---|---|---|---|---|---|---|---|---|
| LerWT | 0.45 | 0.44 | 0.50 | 0.50 | 0.46 | 0.50 | 0.49 | -0.01 | 0.48 | -0.39 | -0.40 | 0.45 | 0.46 | -0.46 | 0.45 | 0.44 | -0.41 | 0.39 | 0.50 | 0.45 |
| *rrd2* | 0.50 | 0.50 | 0.50 | 0.50 | 0.47 | 0.46 | 0.49 | 0.19 | 0.48 | -0.28 | -0.29 | 0.50 | 0.50 | -0.47 | 0.50 | 0.45 | -0.44 | 0.50 | 0.50 | 0.50 |
| *rid4-1* | 0.46 | 0.46 | 0.50 | 0.50 | 0.50 | 0.50 | 0.50 | 0.13 | 0.50 | -0.47 | -0.49 | 0.50 | 0.50 | -0.47 | 0.50 | 0.50 | -0.50 | 0.50 | 0.50 | 0.43 |

| *nad4L* | 41 | 55 | 86 | 95 | 100 | 110 | 131 | 158 | 188 | 197 |
|---|---|---|---|---|---|---|---|---|---|---|
| LerWT | 0.50 | 0.48 | 0.46 | 0.48 | 0.46 | 0.50 | 0.43 | 0.48 | 0.46 | 0.44 |
| *rrd2* | 0.50 | 0.50 | 0.47 | 0.44 | 0.44 | 0.49 | 0.43 | 0.48 | 0.42 | 0.45 |
| *rid4-1* | 0.50 | 0.50 | 0.47 | 0.42 | 0.44 | 0.46 | 0.37 | 0.44 | 0.40 | 0.41 |

| *nad5a* | 155 | 242 | 272 | 358 | 374 | 398 | 494 | 548 | 553 | 598 | 608 | 609 | 629 | 676 | 713 | 725 | 764 | 835 | 863 | 875 | 1275 |
|---|---|---|---|---|---|---|---|---|---|---|---|---|---|---|---|---|---|---|---|---|---|
| LerWT | 0.50 | 0.41 | 0.41 | 0.50 | 0.47 | 0.47 | 0.48 | 0.49 | 0.49 | 0.49 | 0.47 | 0.25 | 0.50 | 0.50 | 0.50 | 0.49 | 0.50 | 0.48 | 0.50 | 0.50 | 0.50 |
| *rrd2* | 0.50 | 0.44 | 0.39 | 0.50 | 0.49 | 0.50 | 0.50 | 0.48 | 0.48 | 0.50 | 0.47 | 0.33 | 0.50 | 0.50 | 0.50 | 0.48 | 0.50 | 0.50 | 0.48 | 0.47 | 0.46 |
| *rid4-1* | 0.50 | 0.48 | 0.39 | 0.50 | 0.48 | 0.50 | 0.50 | 0.49 | 0.50 | 0.50 | 0.50 | 0.34 | 0.50 | 0.50 | 0.50 | 0.48 | 0.48 | 0.48 | 0.50 | 0.50 | 0.50 |

| *nad5c* | 1490 | 1550 | 1580 | 1610 | 1665 | 1731 | 1895 | 1916 | 1918 | 1929 | 1958 |
|---|---|---|---|---|---|---|---|---|---|---|---|
| LerWT | 0.46 | 0.48 | 0.47 | 0.46 | -0.29 | -0.39 | 0.49 | 0.50 | 0.47 | -0.40 | -0.48 |
| *rrd2* | 0.45 | 0.45 | 0.46 | 0.46 | -0.31 | -0.44 | 0.50 | 0.50 | 0.50 | -0.40 | -0.50 |
| *rid4-1* | 0.44 | 0.44 | 0.50 | 0.45 | -0.30 | -0.43 | 0.50 | 0.50 | 0.50 | -0.36 | -0.45 |

| *nad7* | 24 | 38 | 77 | 137 | 200 | 209 | 213 | 244 | 251 | 316 | 335 | 344 | 578 | 679 | 698 | 724 | 734 | 739 | 769 | 789 |
|---|---|---|---|---|---|---|---|---|---|---|---|---|---|---|---|---|---|---|---|---|
| LerWT | 0.44 | 0.50 | 0.50 | 0.50 | 0.50 | 0.50 | 0.14 | 0.50 | 0.50 | 0.50 | 0.50 | 0.50 | 0.50 | 0.50 | 0.50 | 0.50 | 0.50 | 0.50 | 0.50 | -0.40 |
| *rrd2* | 0.50 | 0.50 | 0.50 | 0.50 | 0.50 | 0.50 | 0.15 | 0.50 | 0.50 | 0.50 | 0.50 | 0.50 | 0.50 | 0.50 | 0.50 | 0.50 | 0.50 | 0.50 | 0.50 | -0.40 |
| *rid4-1* | 0.45 | 0.50 | 0.50 | 0.50 | 0.50 | 0.50 | 0.13 | 0.50 | 0.50 | 0.50 | 0.50 | 0.50 | 0.50 | 0.50 | 0.50 | 0.50 | 0.50 | 0.50 | 0.50 | -0.41 |

| | 795 | 926 | 963 | 1050 | 1057 | 1079 | 1082 | 1088 | 1103 | 1124 | 1137 |
|---|---|---|---|---|---|---|---|---|---|---|---|
| LerWT | -0.16 | 0.50 | 0.50 | -0.40 | 0.50 | 0.44 | -0.50 | 0.45 | 0.50 | 0.50 | -0.03 |
| *rrd2* | -0.18 | 0.50 | 0.44 | -0.37 | 0.50 | 0.50 | -0.45 | 0.50 | 0.50 | 0.50 | 0.05 |
| *rid4-1* | -0.20 | 0.50 | 0.46 | -0.37 | 0.50 | 0.50 | -0.46 | 0.50 | 0.50 | 0.50 | 0.04 |

Editing status = U / (C+U) - 0.5

| | | | | | | | | | | |
|---|---|---|---|---|---|---|---|---|---|---|
| -0.5 | -0.4 | -0.3 | -0.2 | -0.1 | 0 | 0.1 | 0.2 | 0.3 | 0.4 | 0.5 |

### Supplementary figure 7 (3 of 3)

*orf240A*

| | 199 |
|---|---|
| LerWT | 0.37 |
| *rrd2* | 0.30 |
| *rid4-1* | 0.42 |

*orfX* (*mttB*)

| | 59 | 97 | 144 | 145 | 161 | 164 | 173 | 177 | 200 | 361 | 364 | 377 | 379 | 387 | 406 | 407 | 409 | 412 | 440 | 474 |
|---|---|---|---|---|---|---|---|---|---|---|---|---|---|---|---|---|---|---|---|---|
| LerWT | 0.06 | -0.24 | -0.21 | 0.33 | 0.15 | -0.01 | -0.07 | -0.45 | -0.49 | 0.12 | -0.25 | -0.46 | 0.26 | -0.42 | -0.09 | 0.03 | 0.12 | -0.09 | -0.07 | -0.30 |
| *rrd2* | -0.04 | -0.26 | -0.20 | 0.18 | 0.06 | -0.08 | -0.12 | -0.38 | -0.41 | 0.00 | -0.22 | -0.40 | 0.15 | -0.39 | -0.20 | -0.04 | 0.05 | -0.16 | -0.13 | -0.29 |
| *rid4-1* | -0.13 | -0.41 | -0.38 | 0.02 | -0.08 | -0.27 | -0.30 | -0.43 | -0.49 | -0.13 | -0.37 | -0.48 | 0.03 | -0.48 | -0.35 | -0.20 | -0.06 | -0.34 | -0.28 | -0.43 |

| | 505 | 530 | 538 | 552 | 581 | 586 | 587 | 643 | 649 | 665 | 666 | 693 | 700 | 705 | 746 |
|---|---|---|---|---|---|---|---|---|---|---|---|---|---|---|---|
| LerWT | 0.17 | -0.15 | 0.02 | -0.45 | -0.22 | -0.44 | -0.06 | -0.14 | 0.03 | 0.01 | -0.35 | -0.21 | 0.01 | -0.28 | -0.10 |
| *rrd2* | 0.02 | -0.15 | -0.04 | -0.37 | -0.26 | -0.34 | -0.05 | -0.13 | -0.03 | -0.03 | -0.28 | -0.17 | -0.02 | -0.23 | -0.08 |
| *rid4-1* | -0.07 | -0.33 | -0.21 | -0.46 | -0.42 | -0.47 | -0.24 | -0.31 | -0.23 | -0.21 | -0.44 | -0.46 | -0.19 | -0.42 | -0.24 |

*rpl2*

| | 212 | 292 | 711 |
|---|---|---|---|
| LerWT | 0.40 | -0.47 | -0.43 |
| *rrd2* | 0.47 | -0.45 | -0.41 |
| *rid4-1* | 0.42 | -0.46 | -0.43 |

*rpl5*

| | 35 | 47 | 58 | 59 | 64 | 92 | 169 | 317 | 329 | 512 |
|---|---|---|---|---|---|---|---|---|---|---|
| LerWT | 0.37 | 0.40 | 0.05 | 0.37 | 0.25 | 0.29 | 0.43 | 0.45 | 0.44 | 0.35 |
| *rrd2* | 0.38 | 0.40 | 0.07 | 0.38 | 0.34 | 0.28 | 0.45 | 0.46 | 0.45 | 0.36 |
| *rid4-1* | 0.31 | 0.37 | -0.50 | -0.48 | 0.28 | 0.18 | 0.43 | 0.44 | 0.45 | 0.28 |
| *rid4-2* | 0.31 | 0.36 | -0.46 | -0.33 | 0.30 | 0.26 | 0.42 | 0.50 | 0.48 | 0.32 |

*rpl16*

| | 34 | 43 | 61 | 209 | 229 | 440 | 506 | 512 |
|---|---|---|---|---|---|---|---|---|
| LerWT | 0.48 | -0.30 | 0.50 | 0.49 | -0.42 | 0.44 | 0.50 | 0.50 |
| *rrd2* | 0.50 | -0.30 | 0.43 | 0.47 | -0.44 | 0.44 | 0.40 | 0.46 |
| *rid4-1* | 0.50 | -0.28 | 0.44 | 0.49 | -0.42 | 0.39 | 0.38 | 0.46 |

*rps3*

| | 64 | 126 | 187 | 509 | 515 | 603 | 887 | 1110 | 1244 | 1260 | 1278 | 1344 | 1351 | 1352 | 1470 | 1532 | 1534 | 1571 | 1580 | 1598 |
|---|---|---|---|---|---|---|---|---|---|---|---|---|---|---|---|---|---|---|---|---|
| LerWT | 0.46 | -0.47 | -0.36 | -0.27 | -0.42 | 0.28 | 0.39 | -0.37 | -0.43 | -0.43 | -0.50 | -0.20 | -0.39 | 0.50 | 0.45 | -0.47 | 0.50 | 0.50 | -0.47 | 0.50 |
| *rrd2* | 0.50 | -0.44 | -0.36 | -0.29 | -0.35 | 0.19 | 0.29 | -0.37 | -0.37 | -0.45 | -0.46 | -0.28 | -0.48 | 0.48 | 0.43 | -0.46 | 0.50 | 0.50 | -0.42 | 0.49 |
| *rid4-1* | 0.47 | -0.44 | -0.38 | -0.30 | -0.36 | 0.16 | 0.27 | -0.36 | -0.36 | -0.45 | -0.44 | -0.49 | -0.45 | 0.46 | 0.43 | -0.46 | 0.48 | 0.50 | -0.41 | 0.50 |
| *rid4-2* | | | | | | | 0.32 | -0.32 | -0.31 | -0.43 | -0.40 | -0.50 | -0.42 | 0.47 | 0.35 | -0.41 | 0.50 | 0.50 | -0.37 | 0.50 |

*rps4*

| | 77 | 88 | 175 | 226 | 229 | 235 | 299 | 308 | 332 | 343 | 377 | 524 | 956 | 967 | 992 | 1042 | 1043 | 1052 | 1057 |
|---|---|---|---|---|---|---|---|---|---|---|---|---|---|---|---|---|---|---|---|
| LerWT | 0.26 | 0.29 | 0.40 | 0.41 | -0.42 | 0.26 | 0.41 | 0.28 | 0.30 | -0.48 | 0.40 | 0.39 | 0.38 | 0.38 | 0.43 | 0.29 | 0.35 | 0.31 | 0.38 |
| *rrd2* | 0.20 | 0.24 | 0.36 | 0.35 | -0.50 | 0.19 | 0.31 | 0.19 | 0.22 | -0.49 | 0.38 | 0.31 | 0.32 | 0.30 | 0.41 | 0.24 | 0.23 | 0.20 | 0.31 |
| *rid4-1* | -0.47 | 0.14 | 0.41 | 0.40 | -0.49 | 0.25 | 0.39 | 0.25 | -0.48 | -0.49 | 0.37 | 0.32 | 0.30 | 0.32 | 0.39 | 0.25 | 0.21 | 0.17 | 0.29 |
| *rid4-2* | -0.36 | 0.09 | 0.35 | 0.36 | -0.50 | 0.14 | 0.31 | 0.15 | -0.42 | -0.49 | 0.34 | 0.24 | | | | | | | |

*rps7*

| | 282 |
|---|---|
| LerWT | -0.19 |
| *rrd2* | -0.31 |
| *rid4-1* | -0.35 |

*rps12*

| | 84 | 104 | 146 | 196 | 221 | 269 | 284 | 285 |
|---|---|---|---|---|---|---|---|---|
| LerWT | -0.04 | 0.14 | 0.14 | 0.06 | 0.05 | 0.08 | 0.07 | -0.13 |
| *rrd2* | -0.08 | 0.07 | 0.08 | 0.03 | 0.01 | 0.02 | 0.02 | -0.18 |
| *rid4-1* | -0.20 | -0.03 | 0.08 | -0.06 | -0.08 | -0.06 | -0.05 | -0.30 |

*rps14* (*φ*)

| | 67 | 162 |
|---|---|---|
| LerWT | -0.38 | 0.37 |
| *rrd2* | -0.35 | 0.38 |
| *rid4-1* | -0.38 | 0.32 |

Editing status = U / (C+U) - 0.5

| -0.5 | -0.4 | -0.3 | -0.2 | -0.1 | 0 | 0.1 | 0.2 | 0.3 | 0.4 | 0.5 |
|---|---|---|---|---|---|---|---|---|---|---|

### Supplementary figure 8

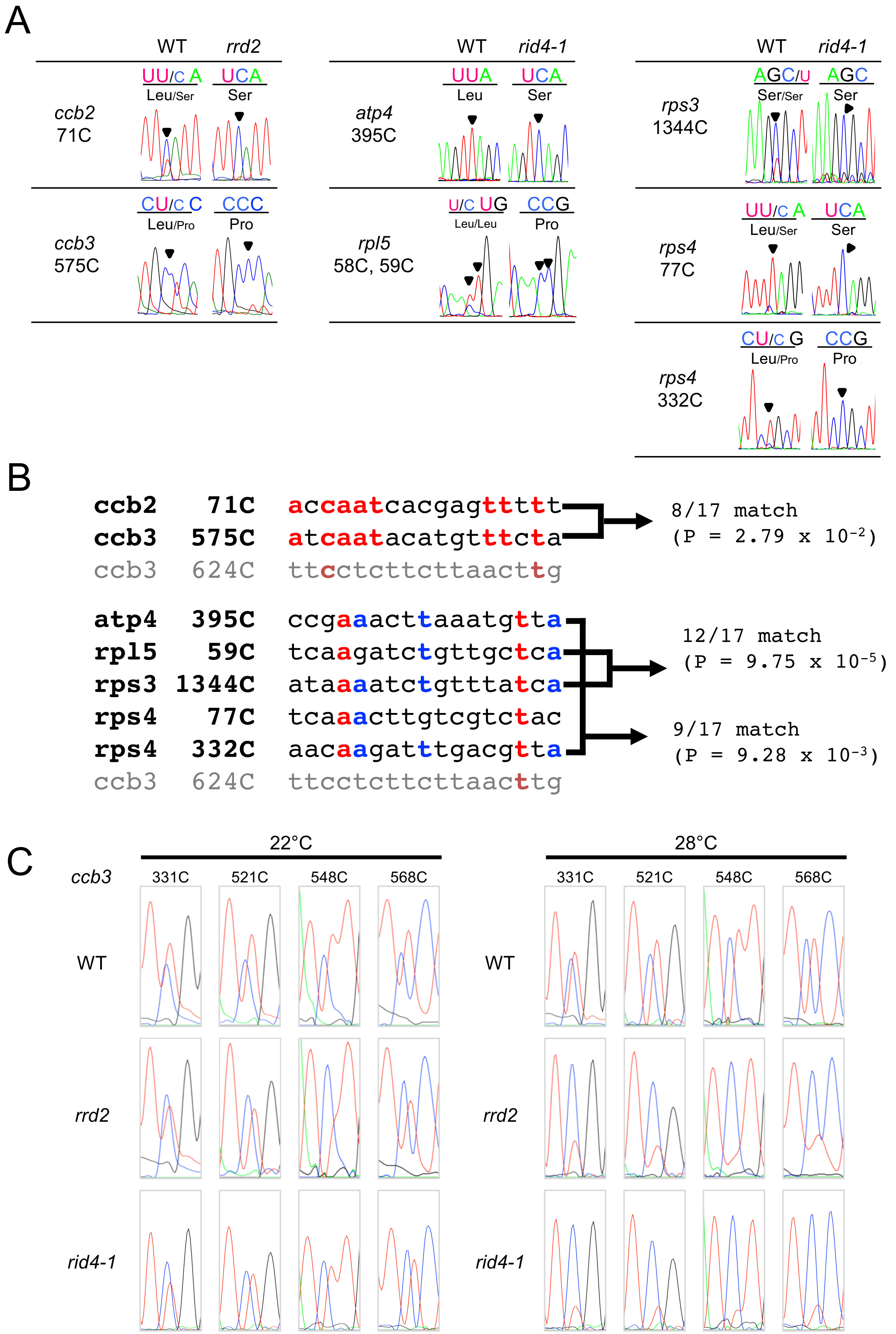

**Supplementary figure 9**

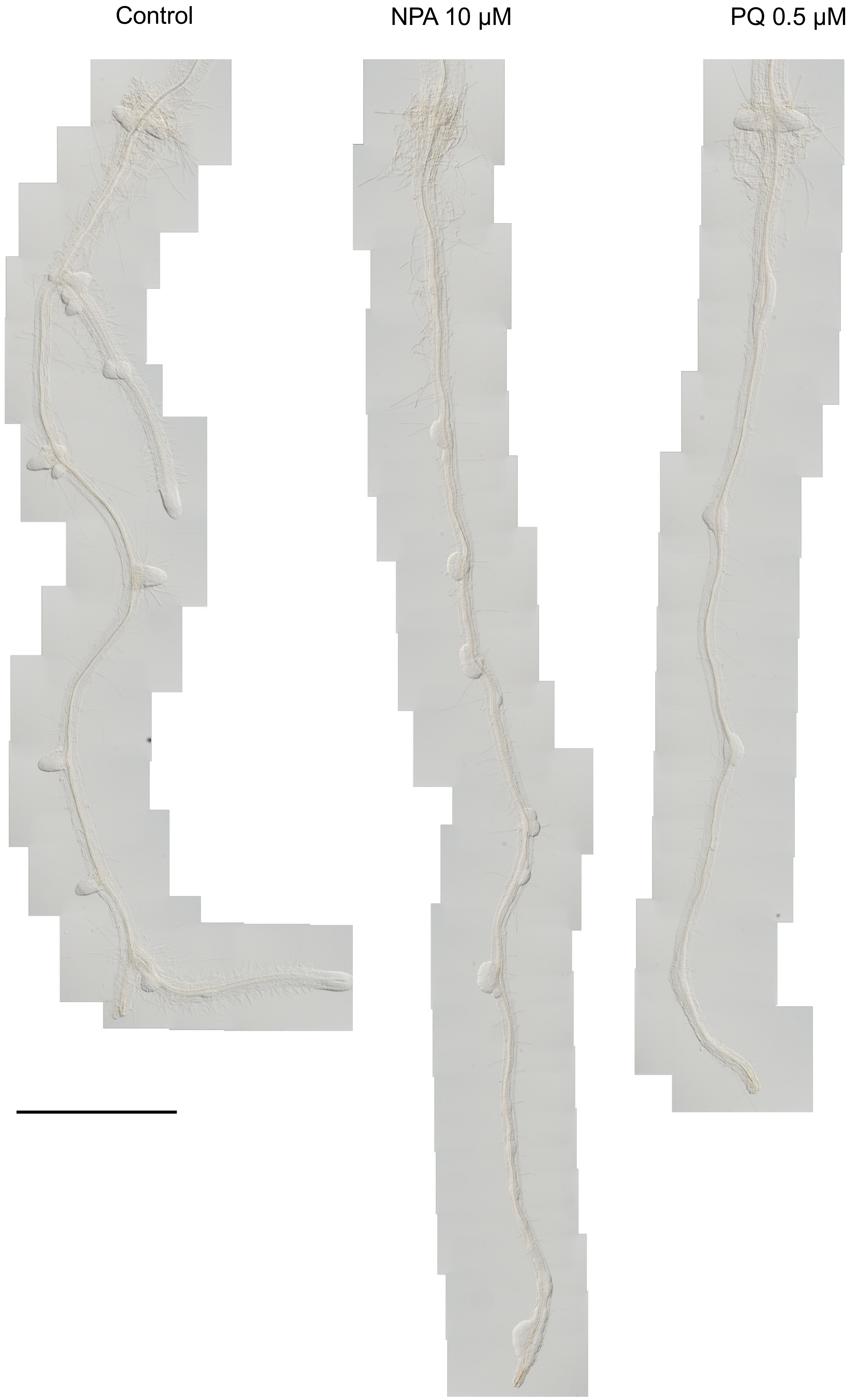

### Supplementary table 1 (1 of 3)

A. Primers for genotyping

| Mutation | Primer name | Primer sequence | Restriction enzyme | Product |
|---|---|---|---|---|
| *rrd1* | dCAPS-rrd1_F | GGAGCTCTGCTATTTTGGTGTTTCG | TaqI | WT = 107 bp, *rrd1* = 24 + 83 bp |
| | dCAPS-rrd1_R | CTGCAACCATCTCTCTTAATGAACC | | |
| *rrd2* | dCAPS-rrd2(NruI)_F | ATGCCTGAGAGAAATGTGGTTTCGCG | NruI | WT = 98 bp, *rrd2* = 24 + 74 bp |
| | dCAPS-rrd2(NruI)_R | GGCATAGCATCAAACACCTGTTTCGC | | |
| *rid4-1* | 2g33680-1F | AACAACGACTCAGTGAGTAC | MvaI | WT = 409 + 732 bp, *rid4* = 1,141 bp |
| | 2g33680-1R | AACAATCGTGTATTCACTGG | | |
| *rid4-2* | dCAPS-rid4-2-F | GTCGAGTCTTCAAAGTTCTACAGCT | XspI | WT = 45 + 68 bp, *rid4-2* = 25 + 20 + 68 bp |
| | dCAPS-rid4-2-R | CTTTACAGTACATGCCCACCAAC | | |
| *ags1-1* | AGS1F5 | CATCTCTATCAAGTTCCTTGATGC | XmnI | ColWT = 153bp, LerWT = 97 + 56bp, *ags1-1*= 131 + 22bp |
| | ags1-1dCAPS3 | GGTTGATCGCAGGAAACGAATTTAT | | |

B. Primers for RACE-PAT assay

| Primer name | Primer sequence | Usage |
|---|---|---|
| T7-Oligo(dT)$_{24}$ | GGCCAGTGAATTGTAATACGACTCACTATAGGGAGGCGGTTTTTTTTTTTTTTTTTTTTTTTT | RT |
| T7-promoter | GTAATACGACTCACTATAGGG | PCR |
| At2g07707_GS-FW1 | AAGTGGCTCACGAGGAATGGAA | PCR, atp8 |
| At2g07727_GS-FW1 | CCTGTGGAGGCTCCATTTGTTACT | PCR, cob |
| cox1_GS-FW1 | GGAGAACTTCCAGCTATCAAGG | PCR, cox1 |
| COX2_GS-FW1 | GGCTTGTATGGCAGGAATACCACT | PCR, cox2 |
| nad6_GS-FW1 | GGGGCTATAGTACTGACTATGC | PCR, nad6 |
| nad9_GS-FW1 | GTGGAAGTACGCTATGATGATCC | PCR, nad9 |
| beta-TUBULIN_FWD | GAGGGAGCCATTGACAACATCTT | PCR, β-TUB4,control |
| beta-TUBULIN_Re | GCGAACAGTTCACAGCTATGTTCA | |

C. Primers for CR-RT PCR

| Primer name | Primer sequence | Usage |
|---|---|---|
| Atcox1-1 | CCCACCAAGAATTTGATCG | Reverse Transcription |
| Atcox1-5'(-176..-196) | TGCTAGGAACGCTAGCTCATG | for PCR |
| Atcox1-3'(+17..+38) | CGACTGCTACTAAGAACCTAAC | for PCR |
| M13 Reverse | CAGGAAACAGCTATGAC | for sequencing |

D. Primers for qRT-PCR

| Primer name | Primer sequence | Gene |
|---|---|---|
| beta-TUBULIN_FWD | GAGGGAGCCATTGACAACATCTT | β-TUB4 |
| beta-TUBULIN_Re | GCGAACAGTTCACAGCTATGTTCA | |
| cox1-qRT-FWD | GGGCTAGACATTGCTCTACATG | *cox1* |
| cox1-qRT-Re | TTCAGGGTATGTCCGACCAAAG | |
| COX2-qRT-FWD | GTGATGCTGTACCTGGTCGTTT | *cox2* |
| COX2-qRT-Re | GGAACAGCTTCTACGACGATAG | |
| nad6-qRT-FWD | GTTTATGCCGGAAAGGTACGAAG | *nad6* |
| nad6-qRT-Re | ACTATAGCCCCAATCATGGCTAC | |
| 2g07727-qRT-FWD | TGGCGGATTGCTTACTACTAGG | *cob* |
| 2g07727-qRT-Re | CAGAATGGGCGTTATGGCAAAG | |
| orf25-qRT-FWD | CTCCAGGCTATTCAGGAAGAA | *atp4* |
| orf25-qRT-Re | CTACTACGGTGCCACAAATTC | |

### Supplementary table 1 (2 of 3)

E. Primers for amplifying cDNA of mitochondrial genes

| Gene | Product size | Forward primer name | Forward primer sequence | Reverse primer name | Reverse primer sequence |
|---|---|---|---|---|---|
| atp1 | 1507 | atp1Fw1 | AGCTGCGGAACTAACGAATCTATTCGA | atp1Rv1 | CTAAATTAAAGCTAAAGCTCTTTCTTTTAAGA |
| atp4 | 489 | atp4Fw1 | GCTGCTATTCTATCTATTTGTGCATTAAGTTCGA | atp4Rv1 | CTCTCGAATGAATTCTACTCTTTC |
| atp6 | 857 | atp6Fw1 | AAGTGTGAGGCATTAACGCA | atp6Rv1 | CCTAATTCCAGACCGGTTAATGCA |
| atp9 | 257 | atp9Fw1 | ATGACAAAGCGTGAGTATAATTCTC | atp9Rv1 | CAGAATACGAATAAGATCAAAAAGGCCA |
| ccb2 | 621 | ccb2Fw | CAGCCTTGAAGTGAATGAATT | ccb2Rv | TTAATCTTGTAAACTAATCGAGACC |
| ccb3 | 740 | ccb3Fw | CTACGCGCAAATTCTCATTGG | ccb3Rv | GAGCGAGTGAACTAGGTTTTGGTA |
| ccb6c | 1350 | ccb6cFw | GGTCCAACTACATAACTTTTTCTT | ccb6cRv | ATTATGAACTCCACGGAACTTTCT |
| ccb6c | 1356 | ccb6cFw2 | ATGGTCCAACTACATAACTT | ccb6cRv2 | TGCAATTATGAACTCCACGGAA |
| ccb6n-A | 511 | ccb6n-A_Fw4 | GCGCCGCTTTATCCTGAAAG | ccb6n-A_Rv4 | CAGGATAGCATTTTGCGCCC |
| ccb6n-A | 561 | ccb6n-A_Fw5 | AAACAACCACCAGCGTTTGG | ccb6n-A_Rv5 | ATCTCGTGCCAGATGCAACA |
| ccb6n-A | 205 | ccb6n-A_Fw1 | ATGTCAATATCAATATATGAATTCTTTCAT | ccb6n-A_Rv7 | AGAAAGGTGCATTAGCGGTT |
| ccb6n_B | 547 | ccb6n-Bfw1 | CGAAGCGTGTCGTTCGTAAT | ccb6n-Brv1 | GAGTGACGTAGATGCACAAGA |
| cob | 1136 | cobFw1 | GACTATAAGGAACCAACGAT | cobRv1 | ATCAGTCTCATCCGTGTAAG |
| cox2 | 760 | cox2Fw2 | ATGATTGTTCTAAAATGGTTATT | cox2Rv1 | TTAATTGATTGGATACCCGAGAACC |
| cox3 | 798 | cox3Fw1 | ATGATTGAATCTCAGAGGCATTC | cox3Rv1 | TCATATACCTCCCCACCAATAG |
| matR | 574 | matR_Fw2 | GGTTGAAGTTTAGACCGCTCAC | matR_Rv2 | ATGCTCTTAACGCTCCCCAC |
| matR | 512 | matR_Fw3 | CTTGCATAGCTCAGGTCGGA | matR_Rv3 | TGTTGGGGAACTCTGCAAGG |
| nad1 | 948 | nad1Fw | CCAGCTGAAATACTTGGAATAAT | nad1Rv | AAAGGTGACTAAAAGACCAGAAAC |
| nad1a | 372 | nad1aFw1 | GTACATAGCTGTTCCAGCTGA | nad1aRv | CCCGCTATAATAATTCCATAAACAC |
| nad1b | 269 | nad1bFw | TCGAAATATGCCTTTCTAGGAG | nad1bRv | TCTACATTATAGCCTGCAACT |
| nad1c | 316 | nad1cFw | TCTTCAATGGGGTCTGCTCT | nad1cRv | AAGGAAGCCATTGAAAGGTG |
| nad2a | 560 | nad2aFw1 | CAGAATTCGTTCGGATCCTC | nad2aRv1 | TATTCCAGAGGAAAATGCAC |
| nad2b | 881 | nad2bFw1 | CTTCGATCAATTAGCCAAGA | nad2bRv1 | GATATGAACTGAGTGCCATT |
| nad3 | 360 | nad3Fw1 | ATGATGTCAGAATTTGCACCAAT | nad3Rv | TTACTCCCGATCCGAAGCAC |
| nad4 | 1482 | nad4Fw4 | GTTAGAACATTTCTGTGAATGCT | nad4Rv1 | TGAAATTTGCCATGTTGCAC |
| nad4L | 278 | nad4LFw1 | TGGATCTTATCAAATATTTCACATTTTCT | nad4LRv1 | TTCTACAGCAATAGTACCTC |
| nad5a | 1425 | nad5aFw3 | TTTGCCCCTGCTCGGTAGTTC | nad5aRv1 | TTGGCCAAGTATCCTACAAAGAGAC |
| nad5c | 497 | nad5cFw2 | CCAATTTTTGGGCCAATTCC | nad5cRv1 | GAAAGACGATCGATTATCTACCCA |
| nad6 | 618 | nad6Fw1 | ATGATACTTTCTGTTTTGTCGAGC | nad6Rv1 | TTAGTAGATCGTGAGTGGGTCA |
| nad7 | 1171 | nad7Fw1 | ATGACGACTAGGAAAAGGCA | nad7Rv1 | CTCCAAACACAATATCTTGAGTACC |
| nad9 | 523 | nad9Fw1 | ACTTTACCCAAGAAATGGGTC | nad9Rv1 | GCTGTTCCCAAGGACTAGC |
| orf240 | 696 | orf240Fw1 | ATGCGTAGTAGCGTTCTAAGATCACT | orf240Rv1 | CAACCAGGACCTTTGGACCTC |
| orfX | 861 | orfXFw1 | AATCCTTCACTTTTAGCTTTGAATTACT | orfXRv1 | TCATTGATAGTTACTTTGCCAGGT |
| rpl16 | 540 | rpl16fw2 | ATGTATTTAACCATAAAATCGATTATGCT | rpl16Rv1 | TTACGACCACTGAACAAACTTGGTTG |
| rpl2 | 1050 | rpl2Fw1 | ATGAGACCAGGGAGAGCAAG | rpl2Rv1 | TCACACAGTGAATAAGGGCTTAGG |
| rpl5 | 558 | rpl5Fw1 | ATGTTTCCACTCAATTTTCATTACG | rpl5Rv1 | TTACTGAGTTTCCCCCTCATC |
| rps12 | 378 | rps12Fw1 | ATGCCCACGTTTAATCAATTGA | rps12rv1 | TCATATCGATTTGGGTTTTTCTGCACCA |
| rps3 | 1665 | rps3Fw2 | ATGGCACGAAAAGGAAATCCGATT | rps3Rv2 | TTCGTACGTTTCGGATATAGCACGTC |
| rps4 | 1089 | rps4Fw1 | ATGTGGCTGCTTAAAAAACTGAT | rps4Rv1 | TTATATGTTTTGGCCACGTCC |
| rps7 | 283 | rps7Fw1 | TAATCAAACTTATGGTTGACGCC | rps7Rv1 | CGATTGGTGGAAGCCAGTCC |
| rps14 (φ) | 234 | pseudorps14Fw1 | ACGTAGATTGCTCGCGGCT | pseudorps14Rv1 | TCGAGATGCTAATCCACGAA |

### Supplementary table 1 (3 of 3)

F. Primers for sequencing cDNA of mitochondrial genes

| Gene | Primer name | Primer sequence | Gene | Primer name | Primer sequence |
|---|---|---|---|---|---|
| atp1 | atp1Fw2 | GCTCAGTTGAAAGCTATGAAACAAG | nad3 | nad3Fw1 | ATGATGTCAGAATTTGCACCAAT |
| atp1 | atp1Rv1 | CTAAATTAAAGCTAAAGCTCTTTCTTTTAAGA | nad3 | nad3Rv | TTACTCCCGATCCGAAGCAC |
| atp4 | atp4Fw1 | GCTGCTATTCTATCTATTTGTGCATTAAGTTCGA | nad4 | nad4Fw1 | AGTGGTCTTATTCTGTGTCCT |
| atp4 | atp4Rv1 | CTCTCGAATGAATTCTACTCTTTC | nad4 | nad4Fw2 | AAGATCAAGGCAGCATATCAGT |
| atp6 | atp6fw2 | GGGCTTGATTTTGGGCGAAG | nad4 | nad4Fw3 | CATTGCTTACTCCTCAGTAGCCCATA |
| atp6 | atp6rv2 | TAGTCCAAGCGAACCCACTT | nad4 | nad4Fw4 | GTTAGAACATTTCTGTGAATGCT |
| atp9 | atp9Fw1 | ATGACAAAGCGTGAGTATAATTCTC | nad4 | nad4Rv1 | TGAAATTTGCCATGTTGCAC |
| atp9 | atp9Rv1 | CAGAATACGAATAAGATCAAAAAGGCCA | nad4 | nad4Rv2 | AGATTTGGCCGTAGA |
| ccb2 | ccb2Fw | CAGCCTTGAAGTGAATGAATT | nad4L | nad4LFw1 | TGGATCTTATCAAATATTTCACATTTTCT |
| ccb2 | ccb2Rv | TTAATCTTGTAAACTAATCGAGACC | nad4L | nad4LRv1 | TTCTACAGCAATAGTACCTC |
| ccb2 | ccb2Fw1 | ATGAGACGACTTTTTCTTGAACTAT | nad5a | nad5aFw1 | GCTCCATGGATCTCATCGGAA |
| ccb2 | ccb2_Fw3 | TGCTTGCCAAAGATCCTACTTC | nad5a | nad5a_Rv2 | GGATTTCCCAACAGCACCAATA |
| ccb2 | ccb2_Fw5 | CGTCGTAACGCCCTTAATGC | nad5c | nad5cFw3 | GCTGGTTCTTCGATCAAGTTT |
| ccb2 | ccb2_Fw7 | TCCAGCAGTGGTTGGAACAG | nad5c | nad5cRv1 | GAAAGACGATCGATTATCTACCCA |
| ccb3 | ccb3Fw | CTACGCGCAAATTCTCATTGG | nad7 | nad7Fw1 | ATGACGACTAGGAAAAGGCA |
| ccb3 | ccb3_Fw3 | TGGGATGCTCGTTTGACCTC | nad7 | nad7Fw2 | ATCTGCCTCTTGGCTTATGT |
| ccb3 | ccb3_Rv4 | GAGGTCAAACGAGCATCCCA | nad7 | nad7Rv2 | CTCGTAATGGTACCTCGCAATTCA |
| ccb6c | ccb6cFw1 | TCAGAGCAAGTCGCCCTATT | nad9 | nad9Fw1 | ACTTTACCCAAGAAATGGGTC |
| ccb6c | ccb6cFw2 | ATGGTCCAACTACATAACTT | nad9 | nad9Rv1 | GCTGTTCCCAAGGACTAGC |
| ccb6c | ccb6cFw3 | TTACATGGACCCACTTTTCATTC | orf240A | orf240Fw1 | ATGCGTAGTAGCGTTCTAAGATCACT |
| ccb6n-A | ccb6n-A_Fw4 | GCGCCGCTTTATCCTGAAAG | orf240A | orf240Rv1 | CAACCAGGACCTTTGGACCTC |
| ccb6n-A | ccb6n-A_Fw5 | AAACAACCACCAGCGTTTGG | orfX | orfXFw1 | AATCCTTCACTTTTAGCTTTGAATTACT |
| ccb6n-A | ccb6n-A_Rv5 | ATCTCGTGCCAGATGCAACA | orfX | orfXFw2 | TACTTCGTGGGTGCAACATCAACA |
| ccb6n-A | ccb6n-A_Rv7 | AGAAAGGTGCATTAGCGGTT | orfX | orfXFw3 | ACGGAGGCCTTTTCGACATT |
| ccb6n-B | ccb6n-Bfw1 | CGAAGCGTGTCGTTCGTAAT | orfX | orfXRv2 | TGTTCTCCATAGCAACTGGGG |
| ccb6n-B | ccb6n-Brv1 | GAGTGACGTAGATGCACAAGA | rpl2 | rpl2Fw1 | ATGAGACCAGGGAGAGCAAG |
| ccb6n-B | ccb6nB_Rv3 | AGCATTTTCTACGGGATCCC | rpl2 | rpl2Rv1 | TCACACAGTGAATAAGGGCTTAGG |
| cob | cobRv1 | AGGTGTGATCAGTCTCATCCGTGT | rpl5 | rpl5Fw1 | ATGTTTCCACTCAATTTTCATTACG |
| cob | cob_Fw4 | AGGAACCAACGATTCTCTCTTCT | rpl5 | rpl5Rv1 | TTACTGAGTTTCCCCCTCATC |
| cox2 | cox2Fw1 | CTCTTATACTCAATGGACGAGGTAG | rpl5 | rpl5 Rv2 | AGACATTCCATGCCCTCGGAG |
| cox2 | cox2Fw2 | ATGATTGTTCTAAAATGGTTATT | rpl16 | rpl16fw2 | ATGTATTTAACCATAAAATCGATTATGCT |
| cox2 | cox2Rv1 | TTAATTGATTGGATACCCGAGAACC | rpl16 | rpl16Rv1 | TTACGACCACTGAACAAACTTGGTTG |
| cox2 | cox2Rv2 | GTCCATTGAGTATAAGAGAGCAAA | rpl16 | rpl16Rv | CGGTAATAGGGAGATCCGCG |
| cox3 | cox3Fw1 | ATGATTGAATCTCAGAGGCATTC | rps3 | rps3Fw3 | CGATCAGGCTCGACGACCG |
| cox3 | cox3Rv1 | TCATATACCTCCCCACCAATAG | rps3 | rps3Fw4 | ACTAAGACCTTAATTGAGTCAGTCA |
| matR | matR_Fw2 | GGTTGAAGTTTAGACCGCTCAC | rps3 | rps3Rv1 | ATATACGGATTCCCTCCACCCCTTT |
| matR | matR_Fw3 | CTTGCATAGCTCAGGTCGGA | rps3 | rps3Rv3 | CCCCGGATTTCGTTTTGCT |
| nad1 | nad1aFw1 | GTACATAGCTGTTCCAGCTGA | rps4 | rps4Fw2 | CCCTATTCTTATCGAAGAGAAGGAA |
| nad1 | nad1aRv | CCCGCTATAATAATTCCATAAACAC | rps4 | rps4Rv1 | TTATATGTTTTGGCCACGTCC |
| nad1 | nad1Fw | CCAGCTGAAATACTTGGAATAAT | rps4 | rps4Rv2 | TTTAGTAGTAGGCGGCATCC |
| nad1 | nad1bFw | TCGAAATATGCCTTTCTAGGAG | rps4 | rps4Rv3 | TGAAGTTCGTTTTGTTCCTCTG |
| nad1 | nad1bRv | TCTACATTATAGCCTGCAACT | rps7 | rps7Fw1 | TAATCAAACTTATGGTTGACGCC |
| nad1 | nad1c_Fw2 | GTTCTTTCTAGGAGGTTGGC | rps7 | rps7Rv1 | CGATTGGTGGAAGCCAGTCC |
| nad1 | nad1cRv | AAGGAAGCCATTGAAAGGTG | rps12 | rps12Fw1 | ATGCCCACGTTTAATCAATTGA |
| nad2 | nad2aFw1 | CAGAATTCGTTCGGATCCTC | rps12 | rps12rv1 | TCATATCGATTTGGGTTTTTCTGCACCA |
| nad2 | nad2aRv2 | CATCGAAATGGTACCAGCCG | rps14 (φ) | pseudorps14Fw1 | ACGTAGATTGCTCGCGGCT |
| nad2 | nad2bFw1 | CTTCGATCAATTAGCCAAGA | rps14 (φ) | pseudorps14Rv1 | TCGAGATGCTAATCCACGAA |
| nad2 | nad2bRv2 | TTGACTTTCGTTTGGGCCA | | | |
